# MYC coordinates tRNA pool expansion during re-entry into proliferation by promoting pre-tRNA maturation

**DOI:** 10.64898/2026.11.12.765319

**Authors:** Min Gao, Chantal Scheepbouwer, Boris Simeonov, Ben G. Ooms, Daniela Salak, Cristina Gómez-Martín, D. Michiel Pegtel, Michael Hackenberg, Alan Gerber

## Abstract

Re-entry into proliferation requires rapid expansion of the translational machinery to support cell growth. How cells replenish abundant, long-lived tRNAs remains unclear. Increased RNA polymerase III activity can lead to accumulation of pre-tRNAs without proportional expansion of mature pools, suggesting that transcription alone may be insufficient. Using differentiated HepaRG cells undergoing re-entry into proliferation, we quantified precursor and mature tRNA dynamics. While the total tRNA pool expanded with cell number, mature tRNA abundance per cell remained stable, accompanied by lower precursor-to-mature ratios and accelerated precursor turnover, consistent with enhanced maturation. MYC depletion impaired precursor turnover and compromised maintenance of mature tRNAs and their processing machinery. Across distinct proliferative transitions, the magnitude of per-cell tRNA accumulation paralleled changes in processing-factor mRNA and protein abundance, with both responses dependent on MYC. These findings identify MYC-dependent coordination of tRNA production and maturation as a mechanism supporting the distinct biosynthetic demands of proliferative transitions.

**Graphical abstract:** 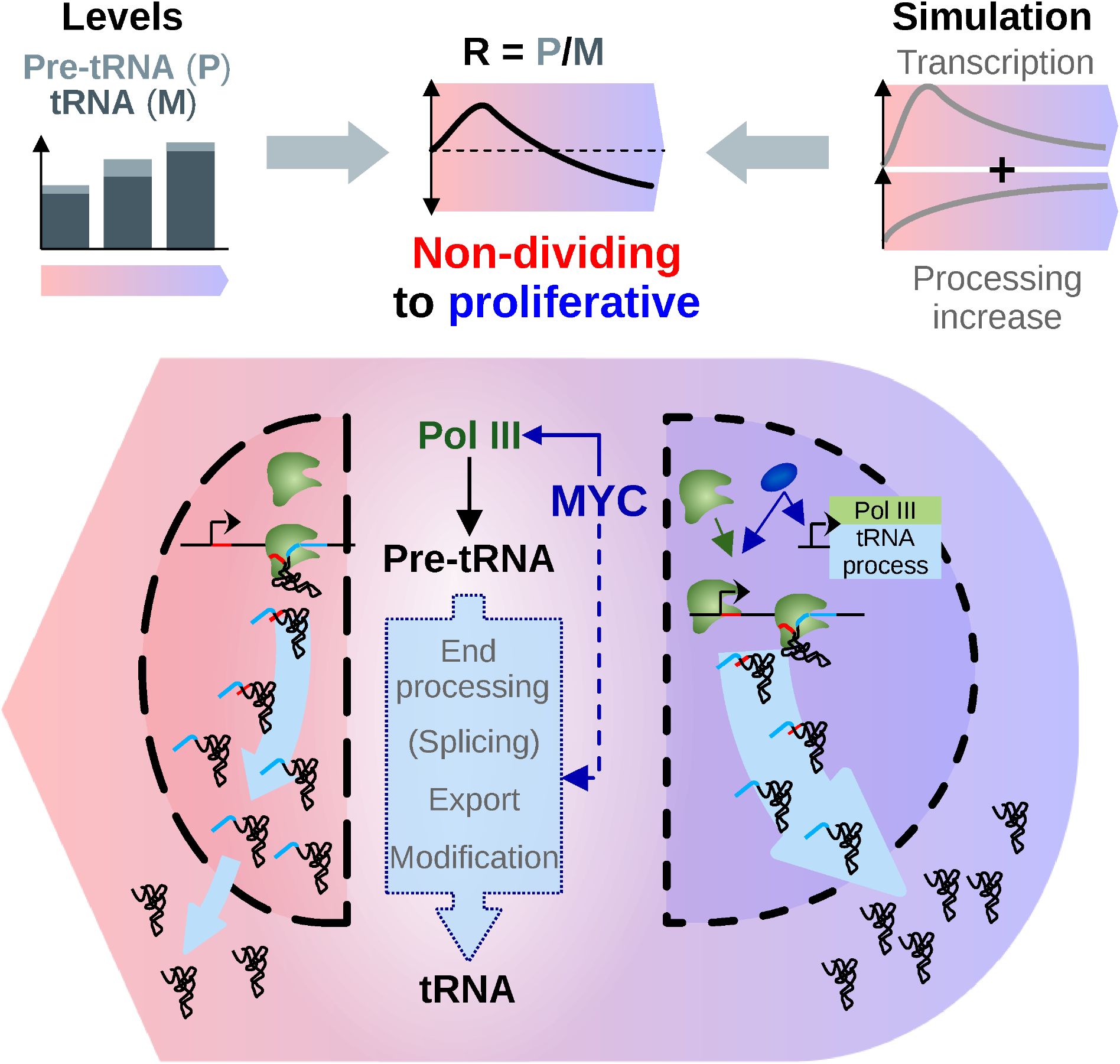

## Introduction

In adult tissues, most somatic cells are non-proliferative. During physiological processes such as tissue repair and maintenance, as well as in pathological contexts including oncogenic transformation, non-dividing cells can re-enter the cell cycle, requiring rapid expansion of their biosynthetic and translational capacity to support growth^1–10^. This creates a regulatory challenge, as core components of the translational apparatus – including ribosomes, transfer RNAs (tRNAs), and associated factors – are both abundant and long-lived, yet must be synthesized at high rates to meet increased demand and counteract dilution caused by cell growth and division. How such abundant and long-lived components can accumulate efficiently during re-entry into proliferation remains unclear.

In particular, mature tRNAs exhibit half-lives on the order of tens to hundreds of hours^11–16^. In non-dividing cells, their long half-lives minimize the need for replacement synthesis. Upon re-entry into proliferation, however, cell growth and division impose a considerable additional requirement for mature tRNA production to maintain these large pools on a per cell basis. In eukaryotes, all cytoplasmic tRNAs are transcribed by RNA polymerase III (Pol III), whose activity is tightly coupled to nutrient- and growth-dependent signaling pathways. At the convergence of these pathways, the conserved repressor MAF1 modulates Pol III transcription according to cellular context. Under growth-promoting conditions, mTOR-mediated phosphorylation of MAF1 relieves repression, leading to increased Pol III occupancy at tRNA genes and upregulation of their expression^17–19^. It remains unclear whether increased Pol III transcription alone is sufficient to rapidly expand functional tRNA pools during cell-cycle re-entry, or whether additional layers of regulation are required.

The relationship between tRNA abundance and translational demand has been extensively discussed in the context of codon usage, where efficient translation is thought to rely on matching between codon bias and the tRNA pool. In this framework, distinct cellular states have been proposed to express specialized tRNA repertoires aligned with specific codon usage programs^20–22^. However, other studies indicate that translational regulation can occur without extensive remodeling of the tRNA pool^23–25^. How these distinct modes of regulation relate to proliferative transitions in mammalian cells remains unclear.

At the level of tRNA production, biochemical and genetic evidence indicate that increased transcription alone has limited capacity to drive productive expansion of the mature tRNA pool. In yeast, *maf1Δ* mutants exhibit elevated Pol III transcription and precursor tRNA levels without proportional increases in mature tRNAs, consistent with saturation of processing, modification, quality control, and export pathways^26–28^. In mice, loss of MAF1 in liver leads to a pronounced, global increase in pre-tRNA synthesis, yet the total mature tRNA pool remains largely unchanged^29–31^. These observations indicate that sustained transcriptional upregulation alone does not drive productive tRNA accumulation, instead resulting in a futile cycle of synthesis and turnover. In contrast, physiological contexts associated with renewed proliferation, such as liver regeneration, display a distinct outcome. Pol III activation during regeneration is more moderate and targeted^32^, yet short-term metabolic labeling experiments demonstrate a substantial increase in mature tRNA synthesis, consistent with increased production rather than altered turnover^33,34^. Notably, regenerated tRNA pools do not exhibit reduced modification levels^33–35^, arguing against relaxation of quality control and indicating that increased tRNA synthesis is coupled to sufficient processing and maturation capacity.

More broadly, MYC coordinates cell cycle progression with increased cellular biosynthetic activity by promoting transcription through all three nuclear RNA polymerases. In particular, MYC directly activates Pol III transcription through interactions with TFIIIB and recruitment of the transcriptional machinery to tRNA and 5S rRNA genes^36,37^. MYC has also been proposed to function as a general transcriptional amplifier, increasing the output of already active genes^38,39^, although the generality of this model remains debated^40^. However, increased Pol III transcription alone would be expected to face the same limitations observed following MAF1 inactivation, where elevated precursor synthesis does not translate into proportional accumulation of mature tRNAs.

Thus, transcriptional output, while necessary, is unlikely to be sufficient for mature tRNA accumulation during cell cycle re-entry. Although further stabilization could, in principle, contribute to tRNA pool expansion, the long half-life of mature tRNAs relative to the cell cycle duration makes this mechanism unlikely to account for substantial accumulation. Instead, tRNA processing – including leader and trailer removal, intron splicing, CCA addition, and nuclear export – emerges as a central regulatory layer^18,41,42^ that may limit productive tRNA accumulation in differentiated cells and must be engaged to support cell-cycle re-entry.

To investigate how this expansion is regulated in human cells, we used differentiated HepaRG cells, a bipotent hepatocyte–biliary-like line which can re-enter proliferation through progenitor-like reprogramming^43–47^. By profiling pre- and mature tRNAs, we assessed how changes in expression and processing contribute to tRNA pool dynamics during the transition from a differentiated to a proliferative state. We show that expansion of the tRNA pool during re-entry into proliferation is not driven by increased transcription alone but is accompanied by coordinated upregulation of pre-tRNA processing. We identify MYC as a central regulator of this response, promoting coordinated expression of genes involved in tRNA maturation and enabling productive accumulation of mature tRNAs upon cell-cycle re-entry. These findings establish regulation of pre-tRNA maturation as a key determinant of translational capacity during transitions to proliferation.

## Results

### Re-entry into proliferation is accompanied by tRNA pool expansion and induction of tRNA biogenesis programs

To determine how mature tRNA pools are expanded during cell cycle re-entry, we used differentiated HepaRG cells, which revert to a progenitor-like state upon withdrawal of differentiation cues and replating at low density. Within 48 h, cells underwent a clear morphological transition from polarized hepatocyte-like clusters to a proliferative epithelial phenotype (**Figure S1A**). This transition was accompanied by a sharp decrease in *CYP3A4* mRNA levels, a hepatocyte marker, and induction of the progenitor and biliary cell marker *KRT19* (**Figure S1B**). Moreover, cells resumed proliferation (doubling time ∼37.8 h; **Figure 1A**), consistent with prior descriptions of this system^43,46,48^.

**Figure 1.**
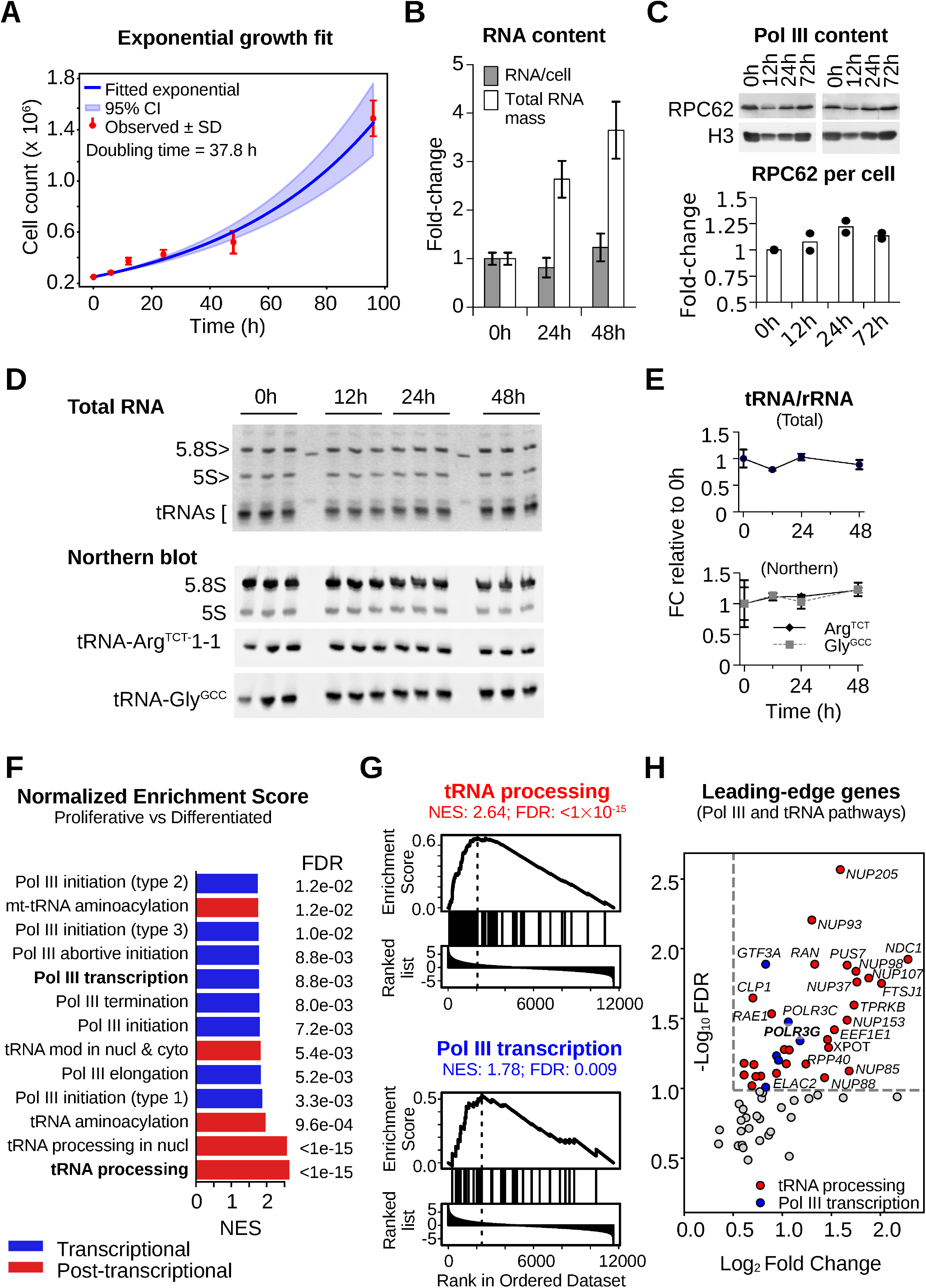
Re-entry into proliferation is accompanied by tRNA pool expansion and induction of tRNA biogenesis programs. **A**) Cell number over time following replating of differentiated HepaRG cells at low density. Data are shown with exponential growth fit (blue), 95% confidence interval (shaded), and observed mean (n = 9). **B**) Total RNA mass per culture and RNA content per cell at 0h, 24h, and 48h. Values are shown as fold-change relative to 0h (n = 9). **C**) Immunoblot analysis of POLR3C (RPC62) and Histone H3, as a proxy for cell number, at indicated time points. Quantification of RPC62 levels normalized to H3 and expressed relative to 0h. Bars indicate mean and points represent individual measurements (n=2). **D**) SYBR Gold-stained denaturing 8% polyacrylamide gel of total RNA (upper part) and northern blotting (lower part) showing 5.8S rRNA, 5S rRNA, and representative cytoplasmic tRNAs (tRNA-ArgTCT-1-1 and tRNA-GlyGCC isodecoders) at indicated time points. **E**) Quantification of **D**) shown as fold-change relative to 0h (n = 3) **F**) Gene set enrichment analysis (GSEA) of differentially expressed genes between proliferative and differentiated states showing terms associated with tRNA or Pol III pathways. Normalized enrichment scores (NES) and false discovery rates (FDR) are shown for selected pathways. Data from ArrayExpress E-MEXP-1082. **G**) Enrichment plots for representative gene sets (tRNA processing and Pol III transcription). **H**) Volcano plot of leading-edge genes contributing to enrichment of Pol III and tRNA-related pathways, plotted by log_2_ fold-change versus -log_10_ FDR. Genes with FDR <0.1 and Log_2_ fold-change >0.5 are colored by Pol III (blue) or tRNA pathways (red). In panels **A**), **B**) and **E**), data represents mean ± SD.

We next asked whether return to proliferation was associated with expansion of the cellular RNA pool. Total RNA mass per culture and RNA per cell were quantified in nine independent experiments at 0 h (differentiated), 24 h, and 48 h post-plating. RNA content per cell was modestly reduced at 24 h (∼1.2-fold decrease, p=0.047) but recovered and slightly increased by 48 h (∼1.2-fold relative to 0 h, p=0.045; **Figure 1B**), consistent with RNA accumulation during re-entry into proliferation. Because changes in cell volume may affect intracellular RNA concentration independently of RNA abundance per cell, we quantified cell size at the same time points (**Figure S1C**). Mean cell volume increased from ∼3.0 pL in differentiated cells to 3.6 pL at 24 h, before returning to 3.0 pL at 48 h. This transient increase followed by a return toward the initial volume is consistent with cell growth and subsequent division, in agreement with the ∼38 h population doubling time (**Figure 1A**). Although cell size distributions differed between conditions, mean cell volume at 48 h was not significantly different from that of differentiated cells (Welch’s test p=0.54). Together with the RNA content measurements, these observations suggest a transient reduction in average cellular RNA concentration at 24 h, followed by recovery and a modest increase at 48 h. Similarly, the overall abundance of Pol III, assessed by POLR3C (RPC62) immunoblotting, largely scaled with Histone H3 levels, a proxy for DNA levels and thus for cell number (**Figure 1C**), indicating that total Pol III levels largely scale with cell number and may be modestly increased on a per-cell basis.

To determine whether tRNA pools expanded proportionally, we quantified total tRNAs relative to 5.8S and 5S rRNAs at 0 h, 12 h, 24 h, and 48 h. Both SYBR Gold staining of total small RNAs and Northern blotting of representative cytoplasmic tRNAs (tRNA-Arg^TCT^-1-1 and tRNA-Gly^GCC^ isodecoders) revealed no significant change in the tRNA-to-rRNA ratio across time points (**Figure 1D** and **1E**). Because rRNAs and tRNAs together account for most cellular RNA, maintenance of this ratio indicates that tRNA levels increase proportionally during cell cycle re-entry. Consistent with this, total RNA mass per culture increased approximately three-to fourfold by 48 h (**Figure 1B**), implying a comparable expansion of the tRNA pool.

We next asked whether this expansion was accompanied by transcriptional changes in pathways related to tRNA biogenesis. Gene set enrichment analysis (GSEA) of differentially expressed genes using a published transcriptomic dataset of differentiated and proliferating HepaRG cells harvested 48 h post-plating^44^ revealed strong enrichment of multiple tRNA-related pathways (**Figure 1F**, **S1D** and **Table S2**). Notably, the most significantly enriched term was tRNA processing (NES > 2.5, FDR < 1×10^-15^), followed by nuclear tRNA processing, and tRNA aminoacylation, along with several pathways associated with Pol III transcription. Representative enrichment plots illustrated increased expression of both tRNA processing and Pol III transcription gene sets **(Figure 1G**), and leading-edge genes contributing to these pathways were predominantly upregulated in proliferative cells (**Figure 1H**), consistent with coordinated activation of tRNA biogenesis. Thus, re-entry into proliferation in HepaRG cells was accompanied by expansion of tRNA pools and coordinated activation of tRNA biogenesis, including components of the processing machinery.

### Pol III isoform switching does not explain tRNA pool expansion during re-entry into proliferation

Given that increased Pol III activity represents the most direct transcriptional mechanism to expand tRNA pools, we first asked whether changes in Pol III composition contribute to tRNA pool expansion during cell cycle re-entry. Pol III composition is regulated through mutually exclusive incorporation of POLR3G (RPC32α) or POLR3GL (RPC32β), which define isoforms associated with proliferative and differentiated states, respectively^49–54^, and have been reported to exhibit context-dependent functions^49,53–55^. *POLR3G* (and *POLR3C*) were among the most strongly induced Pol III-related genes during re-entry (**Figure 1H**).

To test whether Pol III isoform-switching played a role in tRNA pool expansion, we depleted *POLR3G* or *POLR3GL* using pools of four siRNAs in differentiated cells prior to reprogramming and monitored Pol III subunit levels, tRNA accumulation, proliferation rates, and loss of hepatocytic markers during the transition to the proliferative state. In differentiated cells, POLR3G was nearly undetectable at the protein level but increased >16-fold in proliferative cells by 48 h. In contrast, POLR3GL levels slightly declined in proliferative cells over the same time period (**Figure 2A** and **2B**). Depletion of POLR3GL in differentiated cells induced compensatory upregulation of POLR3G, which exceeded 22-fold in proliferative cells. In contrast, POLR3G knockdown prevented its accumulation without inducing POLR3GL expression. Because POLR3G and POLR3GL, together with POLR3C (RPC62) and POLR3F (RPC39), form an essential Pol III initiation subcomplex^56^, we assessed whether changes in isoform composition were accompanied by altered abundance of other Pol III subunits. Neither POLR3G induction nor its depletion altered RPC62 levels relative to RPC53 (**Figure 2A** and **2B**), indicating that these changes were not accompanied by detectable alterations in the relative abundance of the two core subunits.

**Figure 2.**
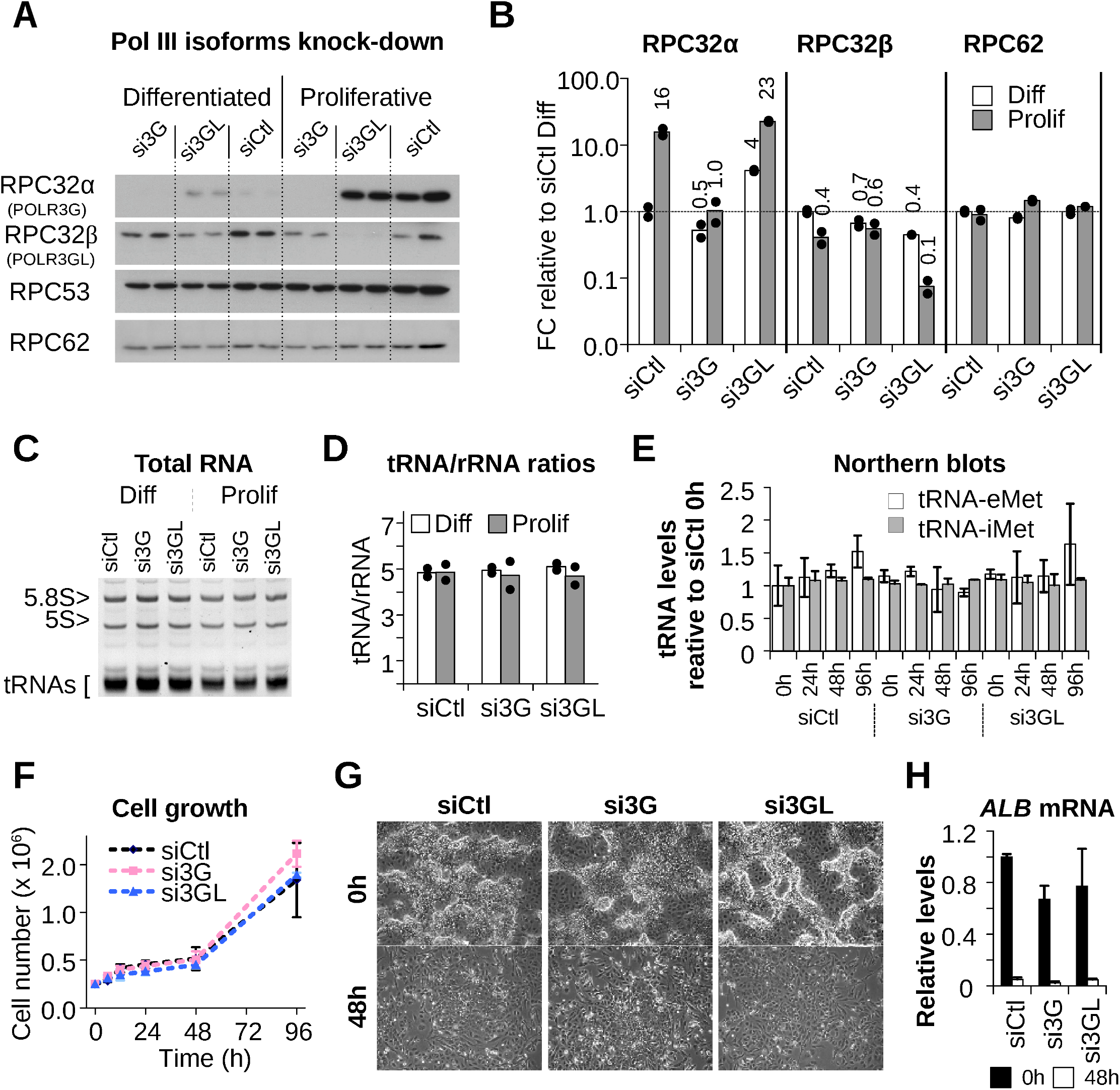
Pol III isoform switching does not explain tRNA pool expansion during re-entry into proliferation. **A**) Immunoblot analysis of POLR3G (RPC32α), POLR3GL (RPC32β), POLR3C (RPC62), and POLR3D (RPC53) in differentiated and proliferative HepaRG cells following siRNA-mediated depletion of *POLR3G* (si3G), *POLR3GL* (si3GL), or control (siCtl). **B**) Quantification of **A**) normalized to RPC53 levels and relative to siCtl-transfected differentiated cells (n = 2). **C**) Representative Ethidium bromide-stained denaturing 6% polyacrylamide gel of total RNA showing 5.8S rRNA, 5S rRNA, and total tRNAs following isoform depletion. **D**) Quantification of tRNA-to-rRNA ratios in differentiated and proliferative cells from two independent experiments as shown in **C**) (n = 2). **E**) Northern blot analysis of two cytoplasmic tRNAs (tRNA-eMet and tRNA-iMet) across time following isoform depletion. Quantification is shown relative to siCtl at 0h; blots are presented in **Figure S2B** (n = 2). **F**) Cell proliferation measured as cell number over time under indicated siRNA conditions (n = 3). **G**) Representative phase-contrast images of differentiated (0h) and proliferative (48h) HepaRG cells under indicated conditions. **H**) RT-qPCR analysis of ALB mRNA levels relative to TBP in differentiated and proliferative cells following isoform depletion (n = 3). In panels **B**) and **D**) points represent individual measurements; bars indicate mean. In panels **E)**, **F)** and **H)**, data represents mean ± SD.

Despite the strong induction of POLR3G in proliferative cells, its depletion had no detectable effect on total RNA content, mature tRNA accumulation, cell proliferation, or de-differentiation (**Figure S2A** and **S2B** and **Figure 2C–2H**). Similarly, POLR3GL depletion did not impair these processes. These results indicate that although POLR3G expression correlates with the proliferative state, Pol III isoform composition is not required for tRNA pool expansion or proliferative reprogramming in HepaRG cells.

### Precursor-to-mature ratios (*R*) distinguish transcriptional from post-transcriptional regulation

If transcriptional machinery composition is not limiting, the key question becomes whether tRNA pool expansion is explained by transcription alone or requires additional post-transcriptional regulation. This is difficult to assess experimentally because pre-tRNAs are short-lived (∼5-15 min half-life^11,57^) and highly sensitive to perturbations, such that procedures required for absolute quantification (e.g., cell counting) are likely to distort precursor dynamics. We therefore analyzed precursor-to-mature ratios (*P*/*M*, denoted *R*), which can be computed directly from relative sequencing data and are inherently insensitive to cell number, global RNA content, and uniform scaling biases. Given the large difference in precursor and mature tRNA lifetimes, *R* captures the relationship between ongoing precursor production and the accumulated mature pool, enabling robust comparison of tRNA dynamics across conditions.

To understand how precursor and mature tRNA abundance are related, we considered a two-compartment kinetic model describing tRNA transcription, precursor maturation, degradation, and growth-associated dilution (**Figure 3A**; Methods and **Supplementary Note S1)**. The balance between mature tRNA production and loss gives *d*ln*(M)*/*dt* = *k_1_*·*R* - *k_degM_ - g*, where *k_1_* is the effective precursor-to-mature conversion rate, *k_degM_* the mature tRNA degradation rate, and *g* the growth-associated dilution rate. Neither transcription nor precursor degradation enters the equation directly, as their effects on mature tRNA production are mediated through precursor abundance, which is captured by *R*. For given rates of mature tRNA accumulation and loss, a reduction in *R* implies a higher precursor-to-mature conversion rate constant.

**Figure 3.**
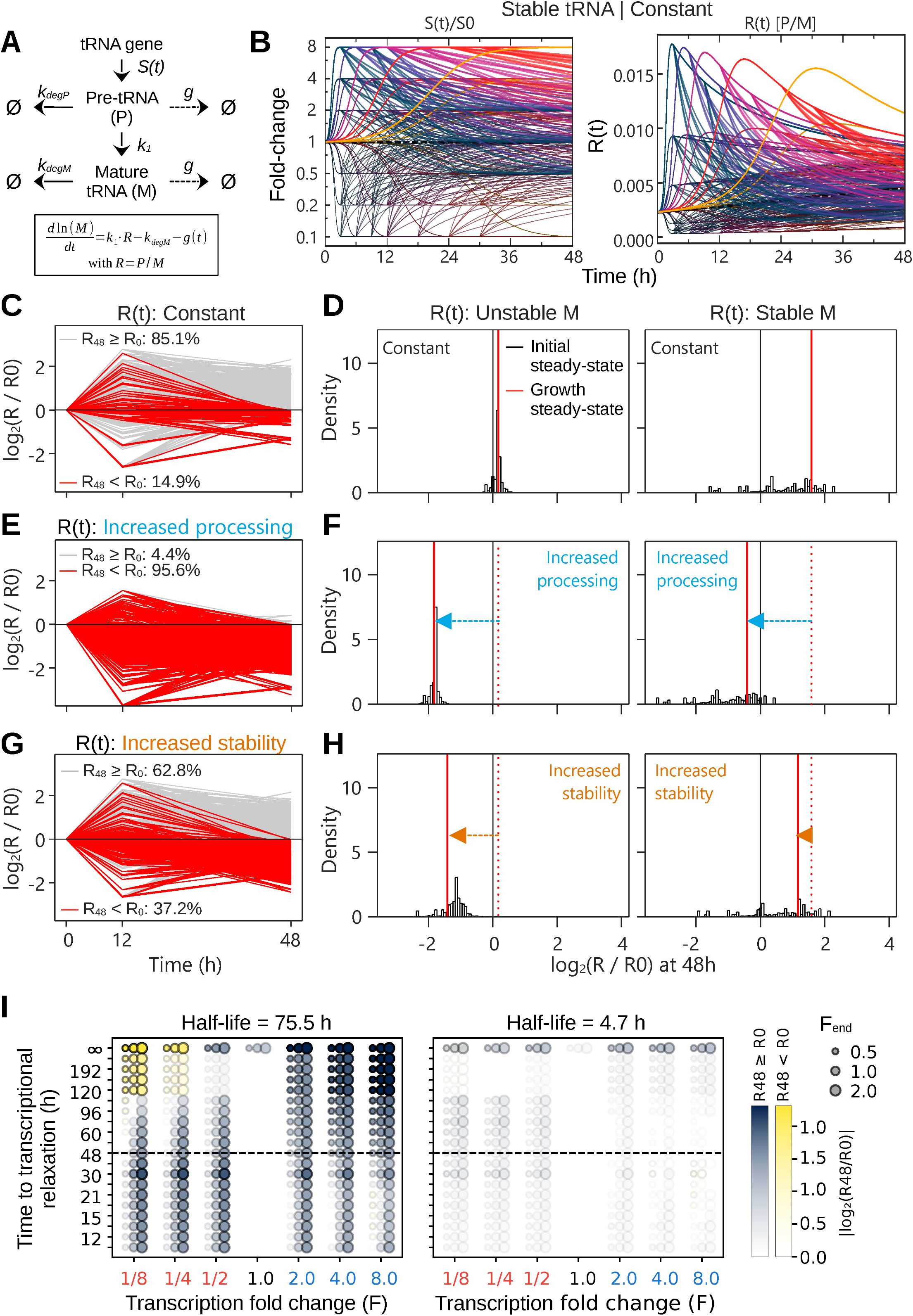
Precursor-to-mature ratios (*R*) distinguish transcriptional from post-transcriptional regulation. **A**) Schematic of the two-compartment model describing precursor (*P*) and mature (*M*) tRNA dynamics, including transcription (*S(t)*), processing (*k_1_*), degradation (*k_degP_*, *k_degM_*), and growth dilution (*g*). Ø denotes loss. The key relationship used in the main text is shown; the full model equations are provided in **Supplementary Note S1. B**) Simulated transcriptional programs (left, fold-change) and corresponding trajectories of precursor-to-mature ratios (*R* = *P*/*M*) over time (right). **C**) Log_2_ fold-change in *R* relative to 0h plotted at 0h, 12h and 48h, and fraction of simulated parameter combinations exhibiting *R_48h_* < *R_0h_* (red) or *R_48h_* ≥ *R_0h_* (gray) under transcriptional modulation alone. **D**) Distributions of log_2_(*R*/*R_0h_*) at 48h under constant conditions for unstable (left) and stable (right) mature tRNAs. The black vertical line indicates *R = R_0h_* (steady state in differentiated cells; 0h baseline), while the red vertical line indicates the expected steady-state shift under proliferation. **E**) Same as **C**) but under increased processing (increased *k_1_*). **F**) Same as **D**) but under increased processing. The dotted red line indicates the expected steady-state shift under constant conditions, and the arrow indicates the direction of the steady-state change induced by increased processing. **G**) Same as **C**) but under increased stability (reduced k_degM_). **H**) Same as **D**) but under increased stability. **I**) Outcomes of transcription-only simulations as a function of peak transcriptional fold change and timing of transcriptional relaxation for short- and long-lived mature tRNAs. Each dot represents a simulated parameter combination. Blue indicates *R_48h_* ≥ *R_0h_*, whereas yellow indicates *R_48h_* < *R_0h_*. Opacity reflects the magnitude of the change in *R*, expressed as | log_2_(*R_48h_*/*R_0h_*)|, with larger changes appearing more opaque. Dot size represents the final transcriptional fold change (*F_end_*). Simulation parameters and definitions are detailed in **Supplementary Note S1**.

Differentiated cells, maintained for extended periods, can be approximated as being at steady state with respect to tRNA abundance, such that *R_diff_* = *k_degM_* / *k_1_, where k_1_*denotes the processing rate constant of a precursor and *k_degM_*the degradation rate constant of the corresponding mature tRNA (see methods section for steady state derivation of *R*). Upon entry into proliferation, growth introduces a dilution term *g*, yielding *R_prolif_*= (*k_degM_ + g*) / *k_1_* at steady state. Because *g* > 0 in proliferating cells, this framework predicts *R_prolif_*> *R_diff_* in the absence of additional regulation. Importantly, transcription does not enter these steady-state expressions, such that *R* is determined solely by processing, decay, and growth. Thus, at steady state, a decrease in *R* during proliferation (*R_prolif_*< *R_diff_*) indicates post-transcriptional modulation. Under growth conditions, processing and decay do not affect *R* symmetrically: for highly stable tRNAs (*g* >> *k_degM_*), modulation of decay has limited impact on *R*, whereas changes in processing directly alter the ratio regardless of tRNA half-life.

However, these predictions are derived at steady state, whereas mature tRNAs are long-lived and proliferating cells may not fully reach equilibrium within the observation window. Nevertheless, growth-induced dilution increases the effective loss of mature tRNAs, shifting *R* toward a higher steady-state value. Thus, growth dilution alone is not expected to reduce *R* below *R_diff_*, making observed reductions suggestive of additional regulation. To assess whether transient transcriptional changes could nevertheless produce such reductions without altering processing or decay, we simulated a two-compartment model (**Figure 3A**) across a broad range of transcriptional programs (**Figure 3B**) varying amplitude, timing, and persistence of induction and repression. Across >1,000 simulated parameter combinations spanning both short- and long-lived tRNAs (half-lives of ∼5 h and ∼76 h), transcriptional modulation combined with growth-induced dilution produced *R_48h_* < *R_0h_* in only a minority of trajectories (∼15% of cases; **Figure 3C**), even though growth steady state was not reached for most long-lived tRNA simulations (**Figure 3D**). In contrast, time-dependent changes in processing or decay readily produced reductions in *R* at 48 h, with ∼96% (processing; **Figure 3E-F**) and ∼37% (decay; Figure **3G-H**) of trajectories satisfying *R_48h_*<*R_0h_*. Notably, decay effects were strongly buffered for long-lived tRNAs, whereas changes in processing robustly decreased *R* across both short- and long-lived tRNAs (compare **Figure 3F** to **3H**). Further examination of the transcription-only simulations revealed that, although ∼15% of parameter combinations produced reductions in *R*, substantial decreases were rare (**Figure 3I**). Only 3.9% of simulations yielded a reduction of at least two-fold, exclusively under eight-fold transcriptional repression in the long-lived tRNA model. Thus, while transcriptional dynamics alone can reduce *R*, pronounced decreases occurred only under a highly restricted set of conditions involving strong transcriptional repression.

However, this framework cannot distinguish whether changes in *R* arise from modulation of processing, decay, or both, nor does it provide the magnitude of these changes. This reflects the fact that precursor and mature tRNAs may differ in their baseline processing and decay rates and may be differentially regulated through either pathway.

These simulations indicate that widespread reductions in *R* are unlikely to arise from transcriptional adaptation alone, even under non-steady-state conditions. We therefore selected three biologically defined time points – 0h (differentiated) as a steady-state reference, 12h (early re-entry), and 48h (proliferative) characterized by exponential growth (**Figure 1A**) and, as shown elsewhere, maximal BrdU incorporation^43^ – to quantify precursor and mature tRNA dynamics during this transition. Hence, these results establish a framework to detect global post-transcriptional contributions to tRNA pool expansion, motivating a direct empirical strategy to quantify precursor and mature tRNA species across this transition.

### Empirical definition of pre-tRNAs from RNA sequencing

To quantify precursor and mature tRNA species within the same sample, we adapted the ALL-tRNAseq^58^ procedure to capture molecules spanning both precursor and mature size ranges. The protocol was modified to recover both capped^59^ and 5′-triphosphorylated precursor transcripts, while reducing abundant short rRNAs to improve library complexity (**Figure S4A, S4B**).

Because existing annotations define only the mature tRNA body, precursor transcripts were identified empirically. Precursor-derived reads were defined as fragments extending beyond annotated mature tRNA boundaries while remaining anchored to tRNA loci, thereby capturing leader and/or trailer sequences (**Figure 4A)**, whereas reads confined to the mature body were excluded. The resulting population exhibited insert sizes larger than annotated mature tRNAs (**Figure 4B**), consistent with enrichment for precursor-containing molecules.

**Figure 4.**
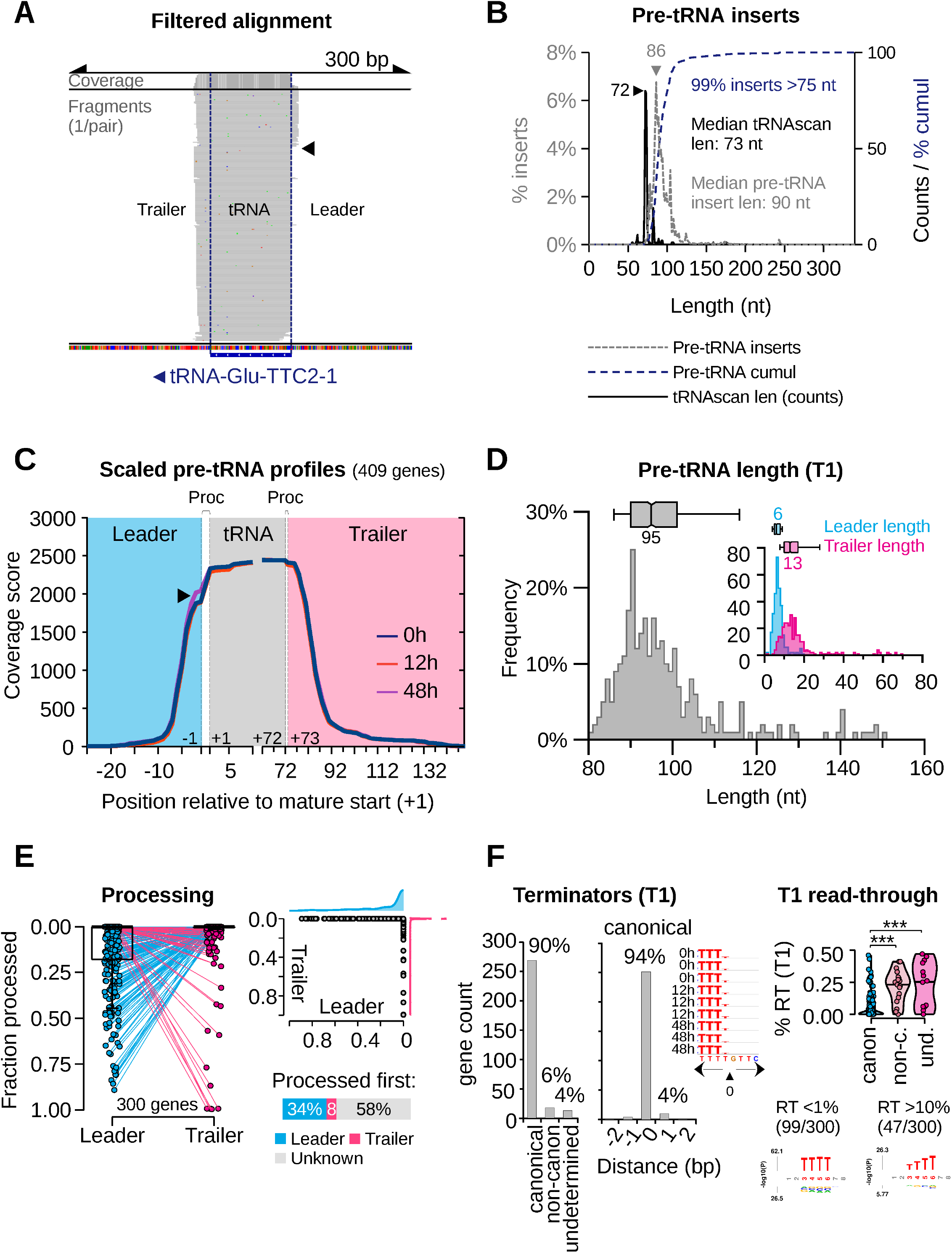
Empirical definition of pre-tRNAs from RNA sequencing. **A**) IGV snapshot of a representative locus showing filtered alignments capturing reads extending beyond annotated mature tRNA boundaries, including leader and trailer regions in one sample. Colored marks within aligned reads indicate nucleotide mismatches relative to the reference sequence. See also **Figure S6A**. **B**) Length distribution of filtered pre-tRNA inserts (gray; left y-axis, percentage) compared to annotated mature tRNA lengths (tRNAscan, black; right y-axis, counts per size). The cumulative distribution of pre-tRNA inserts is shown as a blue dashed line (right y-axis, percentage). Data represents the mean of all sequenced samples across time points. **C**) Normalized and scaled coverage profiles of filtered pre-tRNA inserts across 409 genes at 0h (blue), 12h (red), and 48h (purple), aligned relative to the mature tRNA body (scaled to 72 nt). The arrowhead indicates the only visible difference at the leader cleavage site at 48h. **D**) Distribution and box plot of inferred leader (cyan), trailer (magenta), and full pre-tRNA lengths (grey) based on the operational annotation shown in **Figure S4D. E**) Left: fraction of unprocessed pre-tRNAs per gene, separated by leader or trailer processing. Right (top): fraction of pre-tRNA inserts processed at the leader versus trailer per gene. Right (bottom): inferred fraction of genes showing leader-first, trailer-first, or undetermined processing order. **F**) Left: distribution of transcription termination positions (first drop in coverage downstream of the mature tRNA defined as terminator T1) derived from the operational annotation of pre-tRNAs relative to the end of the T stretch, and quantification of read-through across conditions, including the percentage of canonical (uninterrupted stretch of ≥ 4 Ts), non-canonical (interrupted T stretch), or undetermined (no T stretch) terminators. An inset shows the terminator region at a representative locus across time points, with letter size proportional to sequencing coverage; an arrowhead indicates the operationally defined T1 position. Right (top): percentage of read-through transcription per category (canon, canonical; non-c, non-canonical; und, undetermined). *** p < 1×10^-12^ (ANOVA, Tukey-adjusted p-value). Right (bottom): probability-based logo analysis of terminator sequences at genes showing <1% RT and >10% RT. Statistically significantly enriched positions are shown in red (Bonferroni-adjusted p < 0.01). Letter size corresponds to -log_10_(adjusted p-value).

To characterize precursor structure, we generated scaled coverage profiles in which the mature tRNA body was normalized to a common length and flanking regions aligned relative to this reference (**Figure S4C**). Aggregated profiles revealed consistent upstream and downstream signal corresponding to leader and trailer regions (**Figure 4C**). These profiles were highly reproducible across time points and displayed sharp transitions at the leader-mature boundary, consistent with precise 5′ processing. The amplitude of these transitions decreased at 48 h, hinting at genome-wide modulation of pre-tRNA processing dynamics. In contrast, transitions at trailer regions were less pronounced. Based on these features, abrupt coverage changes were used to define precursor boundaries and infer pre-tRNA lengths across loci (**Figure S4D**). This analysis revealed short, relatively uniform leader sequences (median 6 nt) and more variable trailer lengths (**Figure 4D** and **Table S3**).

We next evaluated whether sequencing-derived profiles recapitulate known features of tRNA maturation. Pre-tRNAs undergo endonucleolytic maturation at both ends, mediated by RNase P (leader removal) and RNase Z (trailer removal). For genes with detectable partially processed intermediates, coverage profiles were strongly dominated by patterns consistent with either leader- or trailer-cleaved species, with little evidence of substantial accumulation of both intermediate types at the same locus. Because removal of both extensions eliminates the precursor-specific features used for their detection, these intermediates provide information about which cleavage occurred first. Accordingly, 34% of genes showed evidence consistent with leader-first processing and 8% with trailer-first processing. This predominance of leader-first processing is consistent with the established role of SSB (La protein), which binds the nascent 3′ oligo(U) terminus and delays trailer processing, thereby favoring 5′ leader cleavage by RNase P before 3′ maturation^41,60,61^. For the remaining 58%, no partially processed intermediates were detected at sufficient levels to determine the processing order (**Figure 4E**). In addition, transcription termination signals, consisting of stretches of four or more thymidines^62,63^, were accurately recovered (**Figure 4F**): 90% of loci contained a canonical ≥4 T tract, and among these, 94% showed termination precisely at the end of the stretch. The remaining loci corresponded to non-canonical^63^ (interrupted) T tracts, and 4% were classified as undetermined (absence of a T stretch). Canonical terminators were associated with minimal read-through, whereas non-canonical or undetermined transcripts exhibited increased downstream signal, indicating that the approach captures terminator-dependent transcriptional behavior.

These results demonstrate that sequencing-derived profiles robustly capture precursor-derived reads and recapitulate key features of tRNA transcription and processing, enabling quantitative analysis of precursor and mature tRNA populations.

### Coordinated global tRNA pool expansion with a pervasive reduction in *R* indicates a major contribution of post-transcriptional regulation

To quantify precursor-to-mature relationships during re-entry into proliferation, we used BBMap ^64,65^ to align reads to a custom reference library containing mature and empirically defined precursor tRNAs. Reads containing precursor-specific features (leader, trailer, or intronic regions) were assigned to empirically defined precursor references, whereas reads lacking such features were assigned to mature tRNAs. We validated this approach by comparison to the established ALL-tRNAseq^58^ pipeline. Quantification of mature cytoplasmic and mitochondrial tRNAs showed near-perfect agreement across all samples (Pearson r ≈ 0.998–0.999; **Figure S5A**), indicating accurate and reproducible estimation of tRNA abundance. Insert size distributions further confirmed recovery of expected tRNA species, with multimodal profiles matching expected length of CCA-containing mitochondrial and type I and II cytoplasmic mature tRNAs (**Figure S5B**), including a substantial fraction of full-length molecules. Having established robust quantification of precursor and mature tRNAs, we next examined how their relative abundance changes during cell cycle re-entry. The fraction of reads mapping to cytoplasmic tRNAs increased progressively from 0 h to 48 h, whereas mitochondrial tRNAs decreased, indicating opposing shifts between these pools (**Figure 5A** and **Table S4**). Principal component analysis separated samples by time point (**Figure 5B**), and pairwise correlations showed high similarity in tRNA expression patterns across conditions and replicates (r ≥ 0.95; **Figure 5C**), consistent with a coordinated temporal transition.

**Figure 5.**
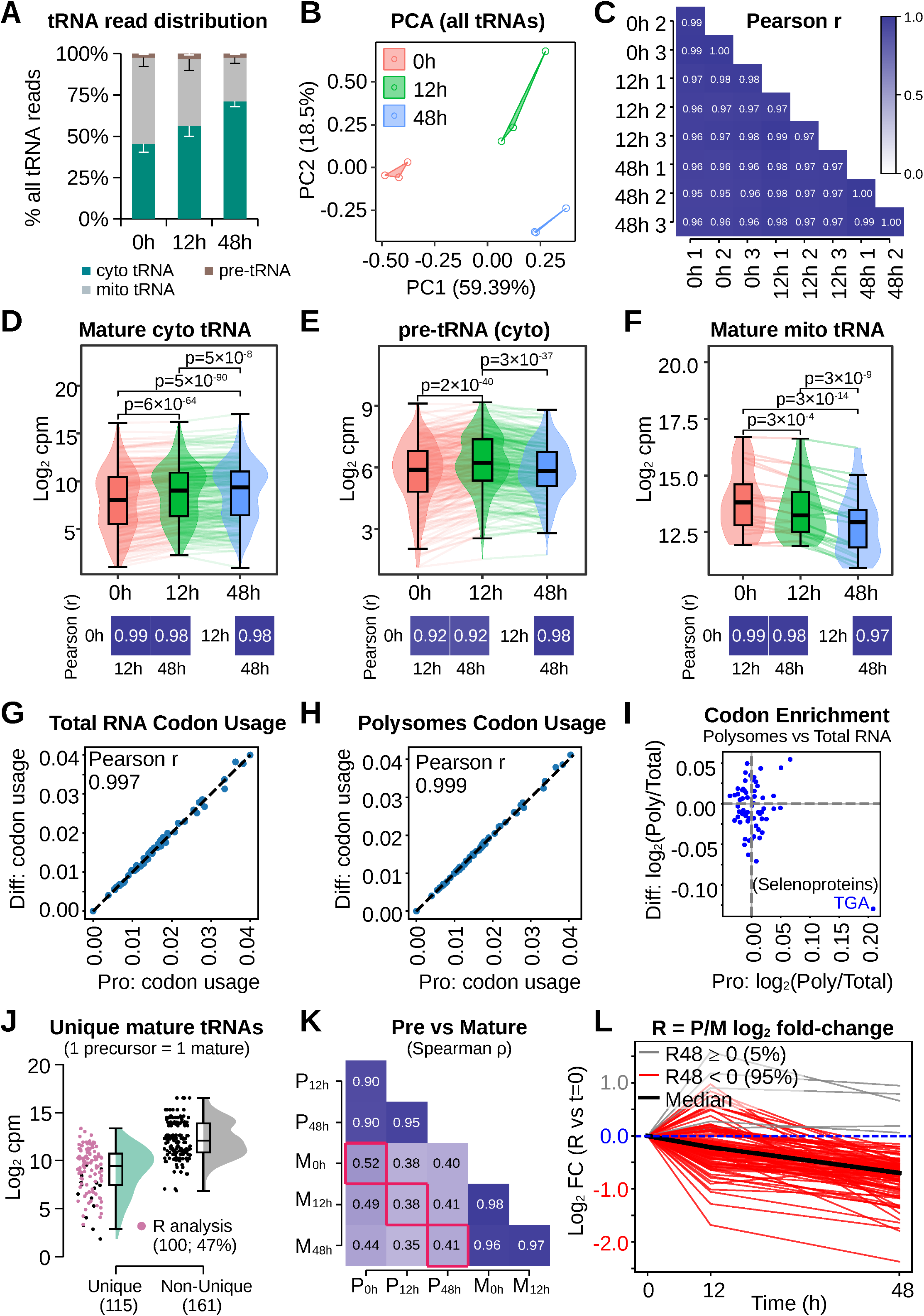
Coordinated global tRNA pool expansion with a pervasive reduction in *R* indicates a major contribution of post-transcriptional regulation. **A)** Fraction of sequencing reads mapping to uniquely assignable cytoplasmic mature tRNAs (260 detected), mitochondrial tRNAs (22 detected), and precursor tRNAs (285 detected) at 0h, 12h, and 48h. Data represents mean ± SD (n = 3). **B)** Principal component analysis of all tRNA species across samples. C) Pairwise Pearson correlation of mature cytosolic tRNA expression across all samples. D)–F) Average expression levels (log_2_ CPM) of mature cytosolic tRNAs (D), precursor tRNAs (E), and mitochondrial tRNAs (F) across time points. Expression changes across time points were assessed using a Friedman test (tRNA species as paired units), followed by Conover post hoc comparisons with Holm-adjusted p-values. Pairwise Pearson correlations between time points for each tRNA group are indicated below the plots to assess preservation of relative expression levels. G) Codon usage derived from total RNA in differentiated and proliferative conditions. Data from ArrayExpress E-MEXP-1082. H) Same as G) but from polysome-associated RNA. I) Log_2_ fold-change in codon demand between polysome-associated and total RNA for each codon in proliferative and differentiated conditions. Statistical significance was assessed by permutation testing with multiple testing correction; no codon showed significant enrichment. J) Expression distribution of uniquely assignable mature tRNAs compared to non-unique species. Pink dots indicate precursor-mature pairs used for downstream analyses (unique only and average expression at each time point ≥ 10 cpm; 100 genes, 47% of expression-filtered precursor-mature pairs). Black dots indicate non-unique expression or average precursor ≤ 10 cpm at any time point. K) Correlation between precursor and mature tRNA levels across time points (Spearman ρ). Red square indicates mature versus precursor at all three time points. L) Log_2_ fold-change of precursor-to-mature ratios (*R* = *P*/*M*) relative to 0h for individual tRNAs.

To distinguish coordinated scaling from compositional changes, we assessed both distributional shifts across genes (Friedman test) and preservation of relative expression across time points (Pearson correlation). Expression levels within each tRNA class remained highly correlated across time points (Pearson r ≥ 0.98 for mature tRNAs), indicating preservation of relative relationships and supporting coordinated scaling rather than compositional reprogramming. Pre-tRNAs showed greater deviation at early time points (r ≈ 0.92 at 12 h), followed by highly coordinated changes between 12 h and 48 h (r ≈ 0.98), with a global shift toward lower relative abundance at 48 h, consistent with transcriptional changes coupled to modulation of precursor processing (**Figure 5D-5F**). In line with this, cytoplasmic and mitochondrial mature tRNAs followed similarly scaled but offset trajectories, whereas precursor tRNAs display non-parallel changes (**Figure S5C**). In addition to preserved tRNA composition, codon usage derived from both total RNA and polysome-associated RNA (reflecting the actively translated transcriptome) remained highly correlated across conditions (Pearson r = 0.997 and 0.999, respectively), with no evidence for codon-specific enrichment when controlling for gene expression levels (**Figure 5G-5I**). These results argue against codon-adaptive remodeling of the tRNA pool during re-entry into proliferation.

Because *R* can only be defined for tRNAs with a one-to-one precursor-mature relationship, subsequent analyses were restricted to uniquely assignable tRNAs. These represented 47% of precursor-mature pairs and spanned a broad range of expression levels (**Figure 5J**), indicating representative coverage of the tRNA pool. Precursor and mature tRNA levels remained highly correlated within each class across time points (r > 0.9), whereas correlations between precursor and mature levels varied, being highest in differentiated cells, reduced at 12 h, and partially restored at 48 h (**Figure 5K** and **S5D**), consistent with transient departure from steady-state relationships. Strikingly, 95 of the 100 expression-filtered uniquely assignable tRNAs (95%) exhibited a reduction in *R* at 48 h relative to the differentiated state (*R_48h_* < *R_0h_*) (**Figure 5L** and **Table S5**). This widespread and coherent decrease is not readily explained by transcriptional modulation alone: although ∼15% of simulated parameter combinations produced reductions in *R*, decreases comparable in magnitude to the experimental median occurred in <4% of cases, all involving strong transcriptional repression (**Figures 3C** and **3I**).

These results demonstrate that re-entry into proliferation is accompanied by a relative increase in cytoplasmic tRNAs, a decrease in mitochondrial tRNAs, and a pervasive reduction in *R* (95%) across tRNA species. Given that total tRNA abundance increases during proliferation (**Figure 1**), these findings indicate coordinated expansion of cytoplasmic tRNAs without major compositional remodeling. As established above, transcriptional modulation alone cannot account for widespread reductions in *R*. The magnitude and coherence of this shift indicate a systematic change in the balance between precursor production and mature tRNA accumulation, supporting a major contribution of post-transcriptional regulation.

### Enhanced pre-tRNA processing accompanies tRNA pool expansion during re-entry into proliferation

To directly assess whether the reduction in *R* reflects changes in precursor processing, we analyzed pre-tRNA processing intermediates using gene-body-scaled coverage profiles. The fraction of unprocessed leader-containing species increased progressively from differentiated to proliferative states (**Figure 6A** and **6B**), indicating a relative depletion of leader-cleaved intermediates during cell cycle re-entry. Direct inspection of genome alignments confirmed that these processing signatures do not arise from fragments lacking overlap with the mature tRNA region, but instead reflect *bona fide* precursor species (**Figure S6A** and **S6B**). In parallel, the proportion of first terminator (T1) read-through transcripts also decreased over time (**Figure 6C**). However, no consistent relationship was observed between read-through levels and the proportion of unprocessed precursors across conditions (**Figure S6C**), arguing against termination efficiency as a primary determinant of precursor processing.

**Figure 6.**
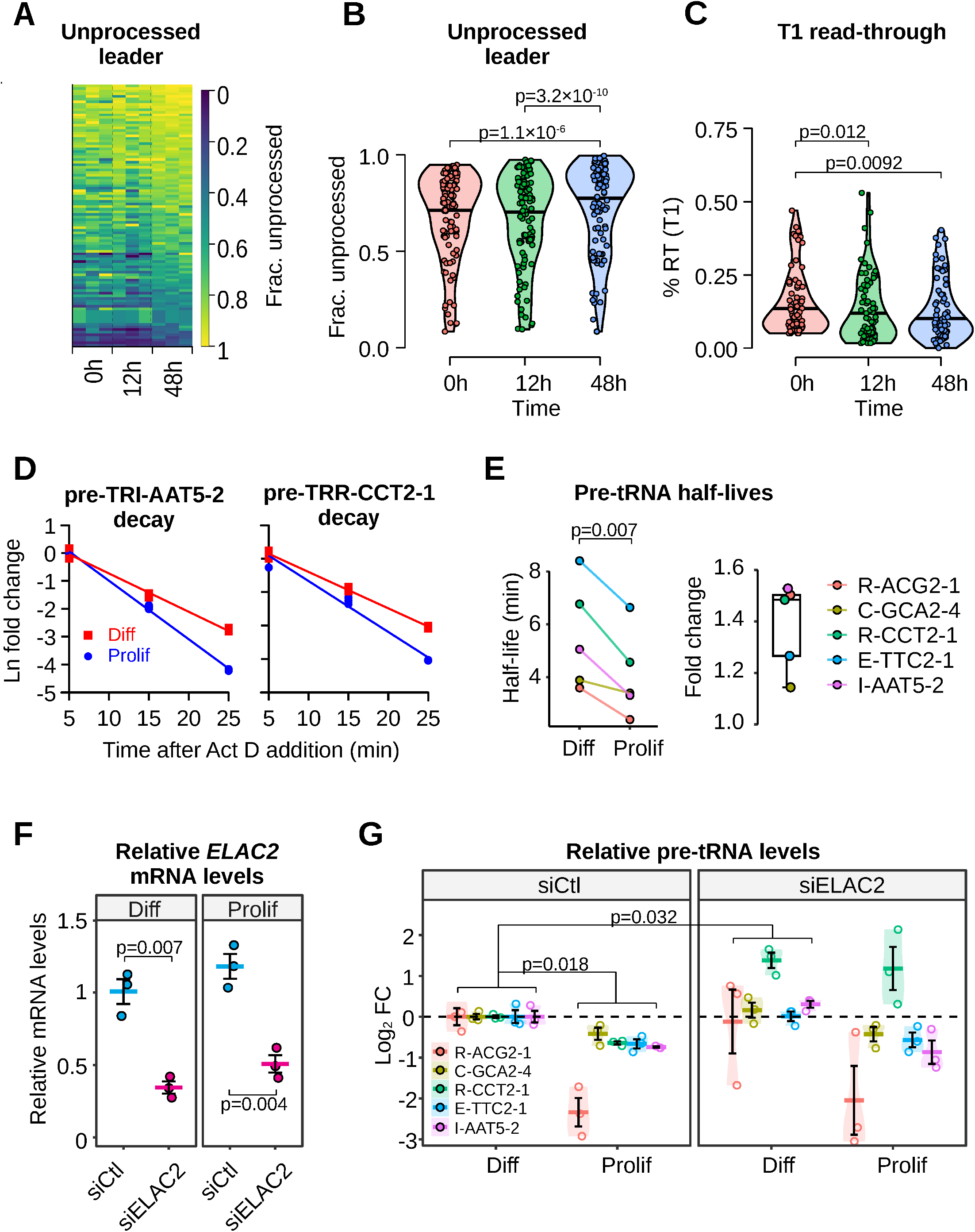
Enhanced pre-tRNA processing accompanies tRNA pool expansion during re-entry into proliferation. **A**) Heatmap of fraction of unprocessed leader-containing pre-tRNAs across time points and replicates. **B**) Quantification of fraction of unprocessed leader-containing pre-tRNAs at 0h, 12h, and 48h. Data represent mean per time point. Only pre-tRNAs showing ≥ 5% processed reads at 0h and SD ≤ 0.2 across replicates were included (101 genes). **C**) Same as **B**) but for T1 read-through levels (≥ 5% at 0h; 73 genes). **D**) Decay kinetics of two pre-tRNAs following transcriptional inhibition assessed by RT-qPCR. Data points represent ln-transformed fold change values normalized to *TBP* and relative to 5 min (beginning of exponential decay phase after initial lag). Two independent biological replicates were analyzed per time point (n = 2). Lines indicate linear regression fits. **E**) Left, estimated half-lives of 5 pre-tRNAs in differentiated and proliferative cells. Right, half-lives fold-change between differentiated and proliferative. Data represent mean (n = 2). Statistical significance was assessed with a paired samples T-test. **F**) RT-qPCR analysis of *ELAC2* mRNA normalized to *TBP* and expressed relative to siCtl-transfected differentiated cells following siRNA-mediated depletion in differentiated and proliferative conditions. Points are individual measurements, dashes indicate mean ± SEM. Statistical significance was assessed with a Welch’s T-test. **G**) Log_2_ fold change of pre-tRNA levels normalized to *TBP* and relative to siCtl in differentiated cells for selected species following *ELAC2* depletion. Points are individual measurements, dashes indicate mean ± SEM. In **B**), **C**) and **G**), statistical significance was assessed using a Friedman test with Conover post hoc comparisons and Holm-adjusted p-values.

To directly examine precursor turnover, we measured pre-tRNA decay kinetics following transcriptional inhibition. Representative pre-tRNAs showed more rapid decay in proliferative cells compared to differentiated cells (**Figure 6D**), resulting in significantly shorter precursor half-lives (**Figure 6E**). While most pre-tRNAs exhibited an approximately 1.5-fold increase in decay rate, others, such as TRC-GCA2-4, showed only minimal changes in turnover. These results indicate that precursor turnover is globally enhanced during re-entry into proliferation, but with gene-specific differences in magnitude. Given the short half-life of pre-tRNAs, their rapid maturation into stable tRNAs, and the expansion of tRNA pools in proliferative cells, these turnover changes are most consistent with altered processing kinetics rather than increased degradation.

Since leader-processed intermediates are relatively enriched in differentiated cells, these observations suggest that 3′ trailer removal may represent a limiting step for a subset of pre-tRNAs in the non-dividing state. To directly test this, we examined the role of ELAC2 (RNase Z), a key enzyme involved in 3′ trailer processing. Depletion of ELAC2 by siRNAs resulted in a significant reduction in mRNA and protein levels in both differentiated and proliferative conditions (**Figure 6F** and **S6D**). In differentiated cells, ELAC2 depletion led to a modest but significant increase in pre-tRNA levels for several tested species. In contrast, ELAC2 depletion had little or no effect on 4 out of 5 tested pre-tRNAs in proliferative cells (**Figure 6G**). However, one pre-tRNA displayed strong sensitivity, with pronounced precursor accumulation upon ELAC2 depletion, indicating that ELAC2-dependent processing can become limiting in a gene-specific manner. Importantly, other pre-tRNAs with similar decay kinetics and fold-changes during cell cycle re-entry were unaffected, demonstrating that distinct subsets of pre-tRNAs are constrained by different steps of the maturation pathway.

Collectively, these observations indicate that re-entry into proliferation is accompanied by a global increase in precursor processing capacity, but that different steps in the maturation pathway impose gene-specific constraints, arguing against a single dominant bottleneck.

### MYC coordinates transcriptional and post-transcriptional components of the tRNA biosynthetic pathway

The observation that pre-tRNA processing is enhanced during re-entry into proliferation, together with the limited or gene-specific impact of perturbing individual components (e.g., POLR3G or ELAC2), suggested that productive tRNA accumulation may depend on coordinated regulation of the tRNA biosynthetic pathway rather than modulation of a single rate-limiting step. We therefore sought to identify upstream regulators of this response and to assess whether it reflects a general feature of proliferative transitions rather than a system-specific effect. To this end, we queried PerturbAtlas^83^, a compendium of published RNA-seq datasets obtained following experimental genetic perturbations, including overexpression, activating mutations, knockdown, and knockout, across diverse cell lines and cellular models. Using the leading-edge genes contributing to enrichment of the Reactome “tRNA processing” pathway in HepaRG cells (**Figure 1F-1H**), we identified candidate positive regulators by selecting perturbations showing enrichment of the tRNA processing gene set following gain of function and depletion following loss of function (**Figure 7A**). This analysis highlighted oncogenic KRAS and MYC among the strongest regulators. Considering the established role of KRAS in activating MAPK signaling and promoting MYC expression and stability^66,67^, these results are consistent with regulation of tRNA biogenesis by mitogenic signaling pathways converging on MYC. These effects were consistently observed across multiple independent datasets and cellular contexts, indicating that this program is a robust and recurrent feature of MYC-associated transcriptional responses. In parallel, transcription factor enrichment analysis across ENCODE datasets identified MYC and its obligate partner MAX as the predominant regulators of these genes, with enrichment scores and odds ratios substantially exceeding those of other transcription factors (**Figure 7B**).

**Figure 7.**
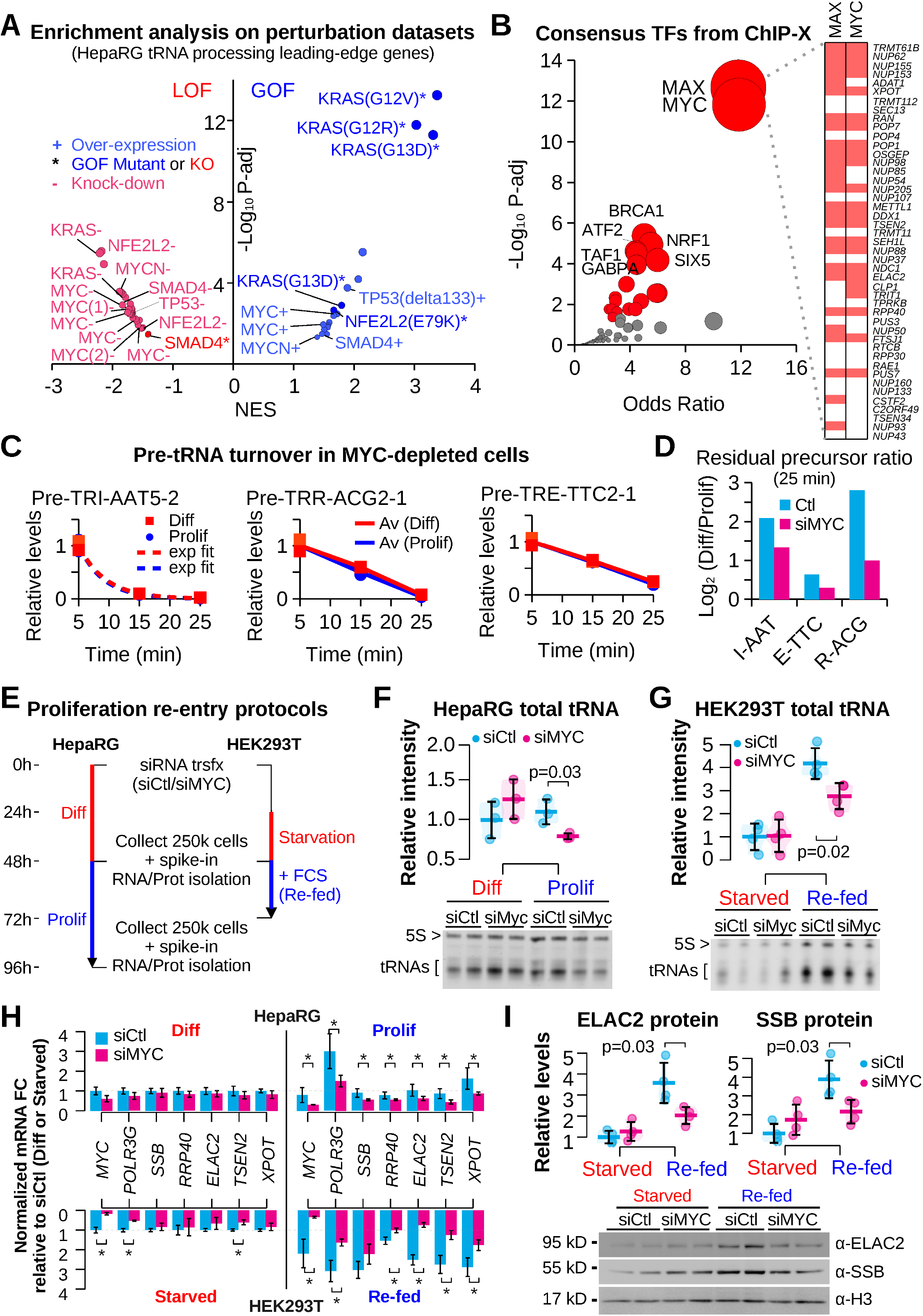
MYC coordinates transcriptional and post-transcriptional components of the tRNA biosynthetic pathway. **A**) PerturbAtlas^83^ enrichment analysis of perturbational RNA-seq datasets using leading-edge genes from the tRNA processing pathway (HepaRG). NES, normalized enrichment score. LOF, loss of function; GOF, gain of function. Only perturbation targets showing reciprocal enrichment following gain of function (NES > 1) and loss of function (NES < -1) were included, selecting for putative positive regulators. **B**) Enrichr^84^ consensus transcription factor enrichment analysis (ENCODE ChIP–seq datasets) for tRNA processing genes. Dot size reflects the Enrichr combined score, integrating enrichment significance and rank deviation; axes indicate odds ratio and –log10 adjusted P value. **C**) Pre-tRNA turnover following transcriptional inhibition in *MYC*-depleted differentiated (Diff) and proliferative (Prolif) HepaRG cells. Representative precursor species (Pre-TRI-AAT5-2, Pre-TRR-ACG2-1, Pre-TRE-TTC2-1) are shown. Precursor abundance was quantified by RT-qPCR normalized to *TBP* and expressed relative to the 5 min time point. Points represent measured values (n = 2); dashed lines indicate exponential fits; solid lines indicate averaged profiles. **D**) Proliferation-associated differences in residual precursor abundance at 25 min following transcriptional inhibition in control (siCtl) and *MYC*-depleted (siMYC) HepaRG cells. Values represent log_2_[(*P*_25_/*P*_5_)Diff/(*P*_25_/*P*_5_)Prolif], where *P*_25_/*P*_5_ is the fraction of precursor remaining at 25 min relative to 5 min. **E**) Experimental overview of proliferation re-entry and matched RNA/protein quantification in HepaRG and HEK293T cells. HepaRG cells were transfected with control or *MYC*-targeting siRNAs in the differentiated state and analyzed before or after re-entry into proliferation. HEK293T cells were transfected with siRNAs, serum-starved, and subsequently stimulated by serum refeeding. For each condition, 250,000 cells were collected and lysed in TRIzol. Exogenous *E. coli* total RNA was added before RNA isolation, and RNA and proteins were recovered from the corresponding aqueous and organic phases of the same biological samples. **F**) Total mature tRNA abundance per cell in differentiated and proliferative HepaRG cells following control or *MYC* knockdown (n=3 for each condition). Representative denaturing RNA gel showing 5S rRNA and mature tRNAs used for quantification of mature tRNA band intensities. Equal volumes of RNA extracts derived from equal cell numbers were loaded without normalization to endogenous rRNA species. Values are expressed relative to control differentiated cells. **G**) Total mature tRNA abundance per cell in serum-starved and serum-refed HEK293T cells following control or *MYC* knockdown, quantified as in F) (n=4 for each condition). Values are expressed relative to control serum-starved cells. **H**) Expression of *MYC* and representative genes involved in Pol III transcription, tRNA processing, and export (*POLR3G, SSB, RPP40, ELAC2, TSEN2, and XPOT*) in differentiated and proliferative HepaRG cells (upper panel) and serum-starved and serum-refed HEK293T cells (lower panel) (n=3 for each condition). RT-qPCR measurements were normalized using exogenous *E. coli* RNA spike-ins (*rho* and *rpoA* transcripts) and expressed relative to control differentiated or serum-starved cells, respectively. HEK293T values are displayed below the horizontal axis for visual separation of the two models; bar lengths represent positive fold changes in expression. Blue and magenta indicate siCtl and siMYC, respectively. **I**) ELAC2 and SSB protein abundance in serum-starved and serum-refed HEK293T cells following control or *MYC* knockdown (n=4 for each condition). Representative immunoblots used for quantification are shown. Proteins were recovered from the organic phase of the same TRIzol samples used for RNA quantification. Histone H3 was used as a proxy for DNA/cell-number loading and to normalize ELAC2 and SSB protein abundance. In F), G), and I), points represent independent biological measurements and horizontal lines indicate mean ± SD. Statistical significance is indicated by the corresponding p-values. In H), asterisks indicate significant differences between the indicated conditions (t-test *p < 0.05).

We next asked whether MYC overexpression induces a similar tRNA biogenesis transcriptional program in an independent cellular model. To this end, we analyzed published transcriptomic data from reprogrammed human hepatocyte (hiHep) organoids overexpressing MYC^68^. This analysis revealed strong enrichment of tRNA-related pathways, including processing, aminoacylation, and multiple Pol III transcription modules (**Figure S7A-S7B** and **Table S6**), closely mirroring the enrichment profile observed in proliferating HepaRG cells. At the gene level, leading-edge genes associated with tRNA processing showed strong directional concordance between HepaRG and MYC-overexpression datasets (**Figure S7C**). This concordance extended across multiple steps of the pathway, including end processing (RNase P/Z), splicing, modification, and export, rather than a single functional module. Genes identified as leading-edge in HepaRG were predominantly regulated in the same direction upon MYC overexpression, even when not classified as leading-edge in that context. In contrast, MYC overexpression induced a broader set of tRNA biogenesis genes, indicating expansion of this program beyond the subset engaged during re-entry into proliferation.

To determine whether MYC contributes to the enhanced processing observed during re-entry into proliferation, we measured pre-tRNA turnover following *MYC* depletion in HepaRG cells. A ∼2-fold reduction in *MYC* mRNA substantially impaired induction of *XPOT* mRNA, a tRNA export factor, in proliferative cells (**Figure S7D**). Despite MYC depletion, cultures continued to expand following replating, with no obvious difference in cell density at 48h. Nevertheless, *MYC* knockdown markedly reduced the proliferation-associated increase in precursor turnover across all tested pre-tRNAs (compare **Figure 6D-6E** to **Figure 7C-7D**). In addition, *MYC* depletion revealed gene-specific alterations in the shape of precursor turnover profiles for a subset of pre-tRNAs, indicating effects beyond canonical leader and trailer processing. Because *MYC* depletion was performed prior to re-entry, both conditions derive from the same perturbed population, enabling direct comparison of turnover dynamics. These findings implicate MYC in the regulation of pre-tRNA maturation during re-entry into proliferation, while suggesting that the effective constraints on processing are distributed and gene-specific rather than governed by a single dominant step.

We next asked whether MYC-dependent regulation of pre-tRNA maturation was accompanied by corresponding changes in mature tRNA abundance and expression of the biosynthetic machinery. To investigate this across distinct proliferative contexts, we compared HepaRG cells undergoing dedifferentiation and re-entry into proliferation with serum-starved HEK293T cells stimulated to resume growth (**Figure 7E**). Importantly, to avoid potential biases introduced by normalization to endogenous RNA species whose abundance may change between cellular states, we collected equal numbers of cells for each condition and isolated RNA and proteins from the same lysates. Exogenous *E. coli* RNA, prepared using LiCl precipitation to preferentially recover larger RNA species and deplete small RNAs, was added to each lysate before extraction to control for RNA recovery and provide an independent reference for mRNA quantification. Equal volumes of the recovered RNA and protein extracts were analyzed, with Histone H3 immunoblotting providing an additional assessment of sample comparability (proxy for DNA content). This approach enabled matched comparisons of tRNA, mRNA, and protein abundance on a per-cell basis.

In HepaRG cells, total mature tRNA abundance per cell remained largely unchanged during re-entry into proliferation, despite ongoing cell division. However, *MYC* depletion significantly reduced tRNA abundance in proliferative cells (**Figure 7F**). In contrast, serum stimulation of HEK293T cells induced an approximately four-fold increase in mature tRNA abundance per cell, which was significantly attenuated by *MYC* depletion (**Figure 7G**). Thus, MYC contributes both to maintaining cellular tRNA pools during proliferative expansion and to their pronounced accumulation following serum stimulation.

We then investigated whether these distinct responses were accompanied by corresponding changes in the expression of the tRNA biosynthetic machinery. In HepaRG cells, expression of most examined processing-factor mRNAs remained relatively stable per cell during re-entry into proliferation, with more pronounced induction of *POLR3G* and *XPOT*. *MYC* depletion reduced the expression of multiple components of the pathway, mainly in proliferative cells (**Figure 7H**). Consistent with these observations, ELAC2 and SSB (La) protein abundance showed relatively modest changes during HepaRG re-entry but were reduced following *MYC* depletion (**Figure S7E**). In HEK293T cells, serum stimulation instead induced substantial increases in the expression of processing-associated genes, accompanied by increased ELAC2 and SSB protein abundance. These responses were markedly attenuated by MYC depletion (**Figure 7H–I**).

To assess the relationship between mature tRNA accumulation and expression of its biosynthetic machinery, we summarized the proliferation-associated changes in total tRNAs, processing-factor mRNAs, and selected processing proteins across both models (**Figure S7F**). In HepaRG cells, all three molecular classes showed relatively limited changes in abundance per cell during re-entry into proliferation, whereas *MYC* depletion compromised their maintenance. In HEK293T cells, the pronounced increase in mature tRNA abundance was accompanied by substantial induction of processing-factor mRNAs and proteins, with *MYC* depletion attenuating these coordinated responses. Thus, the magnitude of tRNA pool expansion paralleled changes in expression of the biosynthetic machinery across distinct proliferative transitions, with both responses dependent on MYC.

Together with the effects of *MYC* depletion on precursor turnover, these findings support a model in which MYC coordinates tRNA production and maturation to meet the distinct biosynthetic demands of proliferative transitions, enabling maintenance or expansion of mature tRNA pools during cell growth and division.

## Discussion

Expansion of the translational machinery is essential for proliferative growth, yet many of its components, including tRNAs, are among the most abundant and stable RNA species in the cell, creating a kinetic challenge for their rapid accumulation during cell-cycle re-entry. In HepaRG cells, we found that the total mature tRNA pool expanded with the growing cell population, while mature tRNA abundance remained largely constant per cell. This maintenance requires sustained tRNA production to compensate for dilution associated with growth and division. Remarkably, expansion of the total mature tRNA pool was accompanied by widespread reductions in precursor-to-mature ratios and accelerated precursor turnover, without proportional increases in precursor abundance. These observations support an important contribution of enhanced pre-tRNA maturation to productive tRNA accumulation.

Our findings further suggest that this response involves coordinated regulation of the tRNA maturation pathway rather than modulation of a single dominant processing step. The heterogeneous consequences of ELAC2 depletion indicate gene-specific constraints on precursor maturation, while transcriptional analyses revealed MYC-associated regulation of factors involved in multiple steps of tRNA biogenesis, including end processing, splicing, modification, and export. Consistent with a broader coordinating role, MYC depletion attenuated proliferation-associated changes in precursor turnover and compromised maintenance or accumulation of mature tRNAs across distinct proliferative transitions. In contrast, depletion of the strongly induced POLR3G-containing Pol III isoform did not impair tRNA accumulation, indicating that Pol III isoform switching is not required for this response. Together, these findings support a model in which MYC coordinates tRNA production and maturation across the biosynthetic pathway to meet the demands of proliferative growth. Given that increased biosynthetic demand characterizes processes such as tissue regeneration and oncogenic transformation, and that dysregulation of tRNA biology is linked to cancer progression^69^, these results suggest that post-transcriptional regulation of tRNA biogenesis may represent an underappreciated component of proliferative adaptation.

The importance of accounting for cell proliferation when interpreting biosynthetic regulation has previously been demonstrated in yeast undergoing stress-induced growth arrest. Lee et al.^70^ showed that substantial reductions in mRNA abundance were not necessarily accompanied by corresponding decreases in protein abundance, partly because reduced cell division diminished the need for protein synthesis to compensate for dilution. Our findings reveal a complementary situation during re-entry into proliferation, where increased dilution creates a requirement for sustained mature tRNA production. Although the underlying kinetic principles are similar, tRNA maturation differs fundamentally from translation because each precursor is consumed to generate a single mature molecule. Thus, regulation of precursor maturation provides a means of adjusting tRNA production to the biosynthetic requirements of cellular growth.

### MYC coordinates the tRNA biosynthetic pathway

MYC is a central regulator of cellular growth, controlling genes involved in ribosome biogenesis, nucleotide synthesis, metabolism, and protein translation^71^. In addition to its role in Pol II-dependent transcription, MYC directly stimulates Pol III activity and promotes rRNA synthesis by Pol I^72^, coordinating the production of multiple components of the translational machinery. Importantly, MYC also regulates post-transcriptional steps of RNA biogenesis. Schlosser et al. ^73^ demonstrated that MYC promotes the maturation of 47S pre-rRNA into mature ribosomal RNAs, while Koh et al.^74^ established that MYC controls the expression of core snRNP assembly factors required for efficient pre-mRNA splicing. Consistent with these observations, kinetic analyses have revealed MYC-dependent changes in pre-mRNA processing dynamics^75^. MYC also regulates aminoacyl-tRNA synthetases, extending its influence to tRNA aminoacylation and translational capacity^76,77^. Our findings extend this regulatory framework to the maturation of tRNA precursors. During re-entry into proliferation, MYC-dependent regulation of tRNA biogenesis is associated not only with the expression of transcriptional machinery but also with changes in precursor turnover and expression of maturation-associated factors. These observations suggest that MYC coordinates precursor synthesis with the downstream maturation capacity required for productive accumulation of mature tRNAs. Thus, rather than acting solely through transcriptional amplification, MYC may coordinate successive stages of RNA biogenesis to meet the biosynthetic demands of proliferating cells.

The importance of MYC in coordinating cellular biosynthetic capacity is also evident in the liver, where MYC overexpression induces ribosomal and translation-associated genes and promotes hepatocyte hypertrophy independently of proliferation^78^. Conversely, genetic studies of liver regeneration have revealed context-dependent requirements for MYC, with restoration of liver mass sometimes occurring despite altered hepatocyte growth and proliferation^79,80^.

An important implication of these findings is that regulation of tRNA biogenesis may depend not only on the biosynthetic demand of a given cellular state, but also on the kinetics of the transition between states. Because mature tRNAs are highly stable, their abundance reflects the balance between production, degradation, and dilution through cell division. During re-entry into proliferation, rapid activation of Pol III transcription could create a transient imbalance between precursor production and maturation capacity, resulting in precursor accumulation without proportional increases in mature tRNAs. As a central integrator of mitogenic signals and cellular biosynthetic programs, MYC is particularly well positioned to coordinate the rapid adjustments in tRNA transcription and maturation required during this transition. Even transient MYC activation may be sufficient to establish the necessary biosynthetic capacity, although whether sustained MYC activity is required once a proliferative state has been established remains unknown.

The upstream signals regulating MYC in this context and the downstream mechanisms through which it enhances pre-tRNA maturation remain to be defined. Perturbation datasets point to a potential role for RAS signaling, as KRAS activation increases expression of tRNA processing-associated genes, whereas its depletion has the opposite effect. RAS-MAPK signaling is known to regulate MYC activity, in part through protein stabilization^66,67^, providing a plausible link between mitogenic stimulation and activation of the tRNA biosynthetic program. MYC mRNA and protein have been reported to undergo transient induction during growth factor-stimulated proliferation in HepaRG cells^81^, consistent with a role in establishing this response. Downstream, MYC may act through transcriptional regulation of processing factors, but the acceleration of precursor turnover despite relatively limited changes in their abundance suggests that additional regulation of processing-factor activity or availability may also contribute.

### Regulation of mature tRNA pools: scaling versus repertoire remodeling

The extent to which tRNA repertoires adapt to changes in translational demand remains debated. While some studies have identified proliferation- and differentiation-associated tRNA programs matching distinct codon usage patterns^20,21^, others have reported remarkable stability of anticodon pools across cellular states^23–25^. These observations suggest that a relatively conserved decoding repertoire may accommodate diverse transcriptional programs without requiring extensive remodeling of tRNA composition. Our findings extend this concept to the transition into proliferation, suggesting that rapid adaptation may primarily require increased production of existing tRNA species rather than redistribution of the anticodon repertoire despite substantial transcript-specific changes in translational output^44^. Such a strategy could allow cells to accommodate renewed growth and division while preserving the decoding capacity already established in the differentiated state.

Importantly, stability of the tRNA repertoire does not necessarily imply unchanged translational regulation. Guimaraes et al.^25^ demonstrated that proliferation-associated mRNAs enriched in rare codons undergo preferential increases in translation efficiency despite little detectable change in the relative abundance of individual tRNAs. Their modeling suggested that a global increase in the availability of translation-ready tRNAs could preferentially relieve elongation constraints at rare codons, providing a mechanism for selective translational regulation without remodeling the tRNA repertoire. Although Guimaraes et al. did not detect a significant increase in bulk tRNA abundance, our observations reveal that the magnitude of mature tRNA accumulation can differ substantially between proliferative transitions, ranging from maintenance of per-cell abundance in HepaRG cells to pronounced accumulation following serum stimulation in HEK293T cells. These differences suggest that global tRNA abundance and functional availability may be regulated according to distinct cellular requirements. Beyond their abundance, tRNA functionality also depends on aminoacylation and post-transcriptional modifications, which can influence decoding efficiency and fidelity. Although our study did not directly assess these properties or their effects on translation, our findings identify regulation of mature tRNA biogenesis as a mechanism supporting tRNA availability during proliferative re-entry. Thus, regulation of tRNA abundance and functional availability may represent distinct but complementary layers of translational adaptation.

In contrast to the coordinated behavior of cytoplasmic tRNAs, the relative representation of mitochondrial tRNAs decreases during re-entry into proliferation, suggesting differential regulation of mitochondrial and cytosolic tRNA pools. In differentiated hepatocyte-like cells, mitochondrial function and oxidative metabolism are prominent, whereas re-entry into proliferation is associated with reduced mitochondrial content and increased biosynthetic demand^82^. This divergence suggests that biosynthetic programs are selectively executed rather than uniformly activated. Taken together, these observations suggest that adaptation during re-entry into proliferation may involve preservation of a core cytoplasmic decoding repertoire coupled to selective adjustments in tRNA functionality and/or compartment-specific biosynthetic requirements.

### Limitations of the study

Several limitations should be considered. First, although precursor-to-mature ratios (*R*), processing intermediate profiles, and precursor turnover measurements provide complementary evidence for enhanced pre-tRNA maturation during proliferative re-entry, they do not directly quantify the flux of precursors into mature tRNAs. Accelerated precursor turnover may reflect increased processing, degradation, or both, and the contributions of individual maturation steps remain unresolved. Direct pulse-labeling approaches are challenging in this system because of the short lifetimes of pre-tRNAs and the long persistence of mature tRNAs. Moreover, although perturbation experiments revealed gene-specific sensitivities, they do not establish which processing steps become rate-limiting for individual pre-tRNAs.

Second, MYC broadly regulates transcription, metabolism, and cell growth, and its depletion may affect tRNA biogenesis both directly and indirectly. While our matched per-cell measurements establish MYC-dependent changes in tRNA abundance and expression of selected components of the biosynthetic machinery, they do not demonstrate that changes in processing-factor expression directly account for altered maturation kinetics. In particular, processing-factor abundance does not necessarily reflect enzymatic activity, and additional mechanisms regulating the activity or availability of existing processing machinery remain to be investigated.

Finally, our experiments specifically address the transition into proliferation, with MYC depletion initiated before cell-cycle re-entry. They therefore do not establish whether sustained MYC activity or enhanced pre-tRNA maturation remains necessary once a proliferative steady state has been established. Temporally controlled perturbations will be required to distinguish regulatory mechanisms involved in establishing proliferative tRNA pools from those required for their subsequent maintenance.

## Material and methods

### Cell culture

HepaRG cells (RRID:CVCL_9720) were cultured as previously described^85^. Briefly, for expansion and de-differentiation, cells were seeded at a density of 2.6 × 10^4^/cm^2^ and grown at 37°C in a humidified atmosphere with 5% CO_2_ in William’s E medium without Glutamine (ThermoFisher Scientific #12551032) supplemented with 10% fetal bovine serum (FBS, ThermoFisher Scientific #A5256701), 1% penicillin/streptomycin, 5 µg/mL recombinant human insulin (Merck #I9278), 50 µM hydrocortisone hemisuccinate (Merck #H2270), and 2 mM GlutaMAX (ThermoFisher Scientific #11574466). For differentiation, cultures were supplemented with 1.8% dimethyl sulfoxide (DMSO; VWR, #23500260) from day 14 post-plating, with medium renewed every 2–3 days. Differentiated cells were washed with PBS and detached by mild trypsinization using 0.25% trypsin–EDTA (ThermoFisher Scientific, #25200056) prior to replating for cell cycle re-entry experiments. All cells were routinely tested and confirmed negative for mycoplasma contamination. HEK293T cells were cultured at 37°C in a humidified atmosphere containing 5% CO_2_ in Dulbecco’s Modified Eagle’s Medium (DMEM, ThermoFisher Scientific #41965) supplemented with 10% FBS and 1% penicillin/streptomycin. Quiescence was induced by serum starvation for 24 h, followed by re-stimulation with 20% FBS for 24 h to induce cell cycle re-entry.

For siRNA-mediated knockdown, cells were transfected with ONTARGET-PLUS siRNA Smart pools (Dharmacon; non-targeting pool [D-001810-10], POLR3G [L-020140-02-0005], POLR3GL [L-014846-02-0005]) or siGENOME siRNA SMARTPools (Dharmacon; non-targeting [D-001206-13-05], MYC [M-003282-07-0005], ELAC2 [M-015640-02-0005]) using Lipofectamine RNAiMAX Transfection Reagent (ThermoFisher Scientific #13778100) according to manufacturer’s instructions. HepaRG cells were transfected in the differentiated state and analyzed 48 h post-transfection or replated and analyzed 48 h after induction of re-entry into proliferation. For HEK293T experiments, siRNA transfections were performed 24 h prior to serum starvation and analyzed either after 24 h starvation (quiescent state) or 24 h following serum re-addition. siRNA sequences are listed in **Table S1.**

### Cell counting and cell-size analysis

For cell growth and RNA content measurements, cell numbers were determined using a hemocytometer, with four technical counts per biological replicate. Total RNA was extracted from the corresponding cultures and quantified spectrophotometrically. RNA content per cell was calculated by dividing the total recovered RNA mass by the number of cells in the culture.

For cell-size measurements, differentiated HepaRG cells were collected at 0h, 24h, and 48h following replating. Cell number and diameter distributions were determined using a CASY cell counter (OMNI Life Science). Measurements were restricted to cells with diameters between 9 and 27 µm to exclude debris and larger aggregates. Size distributions were calculated using 0.5 µm bins and normalized to the total number of cells within this range. Cell volumes were estimated from the measured diameters assuming spherical geometry. Three independent biological replicates were analyzed, each measured in technical duplicate. Technical replicates were combined before calculating biological replicate means and standard deviations.

For matched RNA and protein quantification, cells were detached by trypsinization, washed with PBS, and resuspended in 1 mL PBS. Cell concentrations were determined using the CASY cell counter, following dilution of 50 µL cell suspension in 10 mL counting solution. For each condition, 250,000 cells were collected for subsequent RNA and protein isolation.

### Precursor turnover analysis

Pre-tRNA turnover was assessed in differentiated and proliferative HepaRG cells, including MYC-depleted conditions, by transcriptional arrest with 10 μg/mL actinomycin D (ActD; Thermo Fisher Scientific, #A7592). RNA samples were collected at 5, 15, and 25 min following ActD addition. The 5 min time point was used as reference to account for a short lag prior to the onset of precursor decay. Proliferative cells were obtained by replating siRNA-transfected differentiated cells as described above. Pre-tRNA levels were subsequently quantified by gene-specific RT-qPCR as described below. Half-lives were estimated in control samples only, as precursors in MYC-depleted cells did not uniformly exhibit exponential decay. For control conditions, half-lives were derived from the slope of linear regressions fitted to ln-transformed precursor levels normalized to the 5 min time point, using ln(2)/-slope. To enable comparison across control and MYC-depleted conditions, residual precursor levels were calculated as the ratio of abundance at 25 min relative to 5 min (*P_25_*/*P_5_*). State-dependent differences were summarized as log_2_[(*P_25_*/*P_5_*)*_Diff_*/ (*P_25_*/*P_5_*)*_Prolif_*], representing the state-dependent change in relative amount of precursor remaining at 25 min.

### RNA isolation

Total RNA was prepared using TRIzol Reagent (ThermoFisher Scientific #15596018) as previously described^57^. Cells were directly lysed in TRIzol following removal of the culture medium. After a 5 min incubation at room temperature, samples were centrifuged at 12,500×g for 10 min at 4°C to reduce contamination from insoluble material, including genomic DNA. Phase separation was performed by addition of chloroform (0.2 volumes relative to TRIzol), followed by vigorous mixing for 30 s and centrifugation at 12,500 × g for 15 min at 4°C. The aqueous phase was collected and RNA was precipitated by addition of 0.8 volumes of isopropanol and incubation at room temperature for 10 min, followed by centrifugation at 12,500 × g for 10 min at 4°C. RNA pellets were washed twice with 80% ethanol, air-dried, resuspended in RNase-free water, and incubated for 5 min at 55°C to facilitate dissolution. RNA purity and integrity were assessed spectrophotometrically and by electrophoresis on denaturing polyacrylamide gels which were also used for total tRNA and 5.8S rRNA quantifications.

For matched quantification of RNA and protein abundance per cell, HepaRG and HEK293T cells were collected under the indicated conditions and counted using a CASY cell counter as described above. For each biological sample, 250,000 cells were collected, washed with PBS, and lysed in 500 µL TRIzol. Prior to phase separation, 1 µg of LiCl-precipitated *E. coli* RNA, depleted of low-molecular-weight RNA species including tRNAs, was added to each lysate as an exogenous reference. Following chloroform extraction, RNA was recovered from the aqueous phase and resuspended in 30 µL RNase-free water, while the corresponding organic phase was retained for protein isolation. This procedure enabled matched analysis of mature tRNA abundance, processing-factor mRNA expression, and protein abundance from the same biological samples without relying on endogenous RNA species for normalization across cellular states.

### cDNA generation and qPCR

cDNAs were synthesized from 3 μg of RNA using the Superscript IV First-Strand Synthesis System (ThermoFisher Scientific #18090010) with random hexamers, following the manufacturer’s protocol. Briefly, reactions included an initial primer annealing and extension for 10 min at RT, followed by reverse transcription for 25 min at 50°C, and enzyme inactivation for 10 min at 80°C. SYBR Green qPCR was performed in 25 µL reactions, using 2 µL of cDNAs diluted 1:10 in water and PowerUp SYBR Green Master Mix (ThermoFisher Scientific #A25777). Primers were used at a final concentration of 0.3 μM, and amplification was carried out on an Applied Biosystems 7500 Fast Real-Time PCR System. Unless otherwise indicated, HepaRG RT-qPCR measurements were normalized to *TBP*, previously identified as a stable reference gene in this cell line^86^, whereas HEK293T measurements were normalized to *PGK1*.

For pre-tRNA assays, cDNA synthesis was performed using a mixture of gene-specific primers (GSPs) at a final concentration of 2 μM. Pre-tRNAs were selected to span a wide range of expression levels and to contain sufficiently long trailer sequences to accommodate primer binding. GSPs were designed to anneal within the trailer region, with no complementarity to the mature tRNA sequence for the majority of genes tested and only minimal overlap (≤ 3 nt) in the remaining cases, thereby restricting reverse transcription predominantly to precursor transcripts. The GSP mixture additionally included primers targeting *TBP* and other assay-relevant genes (e.g., *ELAC2* and *MYC*), ensuring that all measurements and normalization were derived from the same reverse transcription reaction. qPCR amplification was performed using a forward primer binding within the tRNA gene body and a reverse primer binding within the trailer region, minimizing overlap with the RT primer where possible.

For the matched per-cell expression analyzes (**Figure 7H**), equal volumes of RNA isolated from same starting cell numbers was analyzed by RT-qPCR as described above except expression levels were normalized using the exogenous *E. coli* transcripts *rho* and *rpoA* to correct for differences in RNA recovery and sample processing. Primer sequences are listed in **Table S1**.

### Northern blotting

Northern blots were performed as previously described^57,58^. Briefly, total RNA was resolved on a denaturing 6% urea-polyacrylamide gel and subsequently transferred onto a HYBOND N+ membrane (ThermoFisher Scientific #RPN2020B) by wet electroblotting for 1 h at 30 V. RNA was UV crosslinked to the membrane (0.24 J and 0.12 J applied sequentially to each side of the membrane) prior to pre-hybridization in ULTRAhyb Oligo Hybridization Buffer (ThermoFisher Scientific #AM8663) for 1 h at 42°C with gentle agitation. Hybridization was performed overnight at 37°C using 50 nM biotinylated oligonucleotide probes. Membranes were washed twice for 10 min at room temperature in 5× SSC containing 0.5% SDS, followed by blocking for 15 min in 2× SSC supplemented with 0.5% SDS and 3% BSA. Membranes were then incubated with streptavidin-HRP (1:40,000; Thermo Fisher Scientific, #N100) in blocking buffer for 30 min at room temperature. Membranes were subsequently washed in 2× SSC containing 1% Triton X-100 and 10% BSA, followed by two washes in 2× SSC for 5 min each. Membranes were briefly rinsed in PBS and signals were detected using SuperSignal West Pico Plus chemiluminescent substrate (Thermo Fisher Scientific, #34580) and imaged on a Uvitec chemiluminescence system. Band intensities were quantified using ImageJ. Probe sequences are listed in **Table S1**. Membranes were stripped by incubation with 0.1% SDS preheated to 100°C, followed by cooling to room temperature; this procedure was repeated twice prior to reprobing.

### Total tRNA quantification per cell

For quantification of mature tRNA abundance per cell, equal volumes of total RNA extracts prepared from identical starting cell numbers were analyzed by denaturing 6% urea-polyacrylamide gel electrophoresis. RNA samples were mixed with an equal volume of 2× loading dye, heated at 98°C for 3 min, and immediately cooled on ice before loading. Following electrophoresis, gels were stained with ethidium bromide and imaged using an OptiGo imaging system (Isogen Life Sciences). Mature tRNA band intensities were quantified using GelAnalyzer 23.1.1. Because RNA samples originated from equal numbers of cells and were recovered in equal volumes, band intensities were compared directly without normalization to endogenous rRNA species. Values were expressed relative to control differentiated HepaRG cells or control serum-starved HEK293T cells, respectively.

### Protein extracts and immunoblotting

Total protein extracts were prepared from the TRIzol organic phase obtained during RNA isolation according to the manufacturer’s protocol. Because TRIzol-derived protein pellets were solubilized directly in 1% SDS, which is incompatible with standard protein quantification assays, pellets were resuspended in a volume of 1% SDS approximately equivalent to the recovered pellet volume. Samples were then mixed with an equal volume of 2× Laemmli sample buffer (240 mM Tris-HCl [pH 6.8], 40% glycerol, 8% SDS, 0.04% bromophenol blue, 5% β-mercaptoethanol), boiled for 5 min at 95 °C, and analyzed by SDS-PAGE and immunoblotting.

For matched RNA and protein analyses, proteins were recovered from the organic phases of the same TRIzol lysates used for RNA quantification, each originating from 250,000 cells. Equal volumes of protein extracts were analyzed by immunoblotting. Histone H3 was used as a proxy for DNA/cell-number loading and to normalize ELAC2 and SSB protein abundance. Relative protein levels were calculated from ELAC2/H3 and SSB/H3 band intensity ratios and expressed relative to control differentiated HepaRG cells or control serum-starved HEK293T cells, respectively.

For immunoblotting of Pol III subunits in siRNA-transfected cells, cells were trypsinized, washed with ice-cold PBS, and lysed in RIPA buffer (25 mM Tris-HCl [pH 7.5], 150 mM NaCl, 1% NP-40, 0.5% sodium deoxycholate, 0.1% SDS) supplemented with 0.5 mM DTT, protease inhibitors (Protease Inhibitor Cocktail Set III, Merck Millipore, 1:100), and phosphatase inhibitors (PhosSTOP, Sigma-Aldrich, 1 tablet per 10 mL) added immediately prior to use. Lysates were mixed with an equal volume of 2× Laemmli sample buffer and analyzed by immunoblotting. Primary antibodies used were rabbit polyclonal anti-RPC32α, anti-RPC32β, anti-RPC53, and anti-RPC62 (in house^50,56^), anti-H3 (#9715, Cell Signaling Technology), anti-ELAC2 (ab251725, Abcam) and mouse monoclonal anti-SSB (mAbcam75927, abcam). HRP-conjugated anti-rabbit IgG secondary antibodies were obtained from GE health care (NA934).

### RNA pre-treatments for sequencing

Decapping (and 5’ triphosphate removal) reactions were performed on 3 µg total RNA using mRNA decapping enzyme (New England Biolabs #M0608) according to the manufacturer’s instructions. Subsequently, 5S and 5.8S rRNA depletion was performed by adapting previously published procedures^87,88^. Briefly, 5S and 5.8S rRNAs were captured by incubating total, decapped RNA with 25µM complementary biotinylated 5S and 5.8S rRNA-oligonucleotides (IDT, see Supplemental Table 1) in a total volume of 100 µL 5x saline-sodium citrate (SSC) buffer (AppliChem #A1396.1000). The reaction mixture was first denatured for 3 min at 90°C, followed by hybridization for 10 min at 65°C. Streptavidin-coated magnetic beads (ThermoFisher Scientific #65801D) were washed 3 times in Bind and Wash buffer (5 mM Tris-HCl (pH 7.5), 0.5 mM EDTA, and 1 M NaCl), followed by one wash in 5× SSC buffer. Hybridized RNA samples were then immobilized onto 25 µL primed streptavidin-coated magnetic beads (ThermoFisher Scientific #65801D) to a volume of 100 µL at 600 rpm for 30 min at RT. 5S and 5.8S rRNA-depleted RNA supernatants were then recovered following immobilization of the beads on Magnetic racks 2 min. Depleted rRNA samples were then subsequently cleaned up and concentrated using an Oligo Clean&Concentrator-5 kit (ZYMO Research #D4060). Next, tRNA deacylation was performed by incubating total RNA in 0.1 M Tris-HCL (pH9.0) for 30 min at 37°C, followed by dephosphorylation with 10U T4 Polynucleotide Kinase (PNK; New England Biolabs #M0201L) for 30 min at 37°C, and final purification using an Oligo Clean&Concentrator-5 kit (ZYMO Research #D4060). Demethylation reactions were performed as previously described^58^. Briefly, 2 µg total RNA was treated with 8.5 µM AlkB WT enzyme and 10.5 µM AlkB-D135S mutant enzyme, 10 mM KCl, 2 mM MgCl_2_, 283 µM freshly prepared (NH_4_)_2_Fe(SO_4_)_2⋅_ 6H_2_O, 0.3 mM 2-ketoglutarate, 2 mM freshly made L-ascorbic acid, 40U RNase Inhibitor, and 50 mM MES buffer pH 5.0 (Fisher Scientific #15474529) in a total volume of 40 µL. Reactions were incubated for 2 h at 25°C and quenched with 5 mM EDTA (Ambion #AM9260). RNA was recovered by an Oligo Clean&Concentrator-5 kit (ZYMO Research #D4060) and eluted in 6 µL Nuclease-free water (Ambion #AM9937).

### Expression and purification of MarathonRT

Protocols for expression and purification of MarathonRT were a kind gift from Anna Pyle and were adapted from different studies^89–91^. Briefly, the plasmid pET-6xHis-SUMO-MarathonRT was a gift from Anna Pyle (Addgene plasmid # 109029; http://n2t.net/addgene: 109029; RRID:Addgene_109029) and was overexpressed in *E. coli* Rosetta II (DE3) cells (Millipore). Cells were cultured in 2 L of LB medium supplemented with 50 µg/mL kanamycin and 17 µg/mL chloramphenicol at 37°C to an OD600 of 2.0 to 2.5. The temperature was then lowered to 16°C, and IPTG was added to a final concentration of 0.5 mM to induce expression of MarathonRT. After 18 h, cells were harvested by centrifugation at 2,000 × g for 10 min. Cells were pelleted and resuspended in lysis buffer (25 mM Na-HEPES [pH 7.5], 1 M NaCl, 10% glycerol, and 2 mM β-mercaptoethanol). Cells were lysed by sonication (8 times 10 sec, with 1 min rest) and then centrifuged at 13,000 × g for 30 min at 4°C to remove cell debris. The supernatant was then incubated with 2 mL of pre-equilibrated TALON Metal Affinity resin (TAKARA #CL635502), and the mixture was incubated at 4°C for 1 h. After incubation, the lysate-resin mixture was loaded onto a 30 mL gravity column. The TALON Metal Affinity resin was washed sequentially with 20 mL of lysis buffer, 20 mL of an imidazole-containing buffer A (25 mM Na-HEPES (pH 7.5), 500 mM NaCl, 20 mM imidazole, 10% glycerol, and 2 mM β-mercaptoethanol), and 20 mL of an imidazole-containing buffer B (25 mM Na-HEPES (pH 7.5), 500 mM NaCl, 30 mM imidazole, 10% glycerol, and 2 mM β-mercaptoethanol). To elute the protein, 20 mL of elution buffer (25 mM Na-HEPES (pH 7.5), 300 mM NaCl, 300 mM imidazole, 10% glycerol, and 2 mM β-mercaptoethanol) was applied. MarathonRT, expressed as a fusion protein (N-His6-SUMO-MarathonRT), was then treated with recombinant N-His6-Ulp1 protease (Merck #SAE0067) at 4°C for 1 h to cleave off the N-His6-SUMO tag, yielding the untagged native protein. After protease digestion, the mixture was loaded onto a 5 mL HiTrap SP HP cation-exchange column (GE Healthcare Life Sciences #29051324) equilibrated with 100% low salt buffer (25 mM K-HEPES (pH 7.5), 300 mM KCl, 10% glycerol, and 1 mM DTT). The protein was eluted using a 100% high salt buffer (25 mM K-HEPES (pH 7.5), 2 M KCl, 10% glycerol, and 1 mM DTT) over 20 column volumes (100 mL). Peak fractions were pooled, concentrated to 5 mL, and further purified in 100% low salt buffer by size-exclusion chromatography using a HiLoad 16/600 Superdex 200 column (GE Healthcare Life Sciences #28990944) using an Ultimate 3000 High Performance Liquid Chromatography (HPLC) system (Dionex, Sunnyvale, CA, USA). Protein aggregates were removed by centrifugation on a centrifugal filter unit with 0.22 µm Durapore PVDF membrane (Millipore Sigma #UFC30GV25) and 0.5 mL sample was injected. Detection was carried out at 280 nm and collected fraction size was 3 mL. Collected fractions were pooled, concentrated and then stored at -80°C. All protein purification steps were performed at 4°C. Protein concentration was measured by Pierce BCA protein assay kit (Thermo Fisher #23227). Yield and protein quality were further assessed by SDS-PAGE and Coomassie staining.

### RNA-seq library preparation

Treated RNA was used for each sample. For sequencing libraries, a 5’ phosphorylated 3’ end adapter with 4 randomized nucleotides at the 5’ end and a 3’ blocking group (3C Spacer; 3SpC3, IDT) was adenylated using Mth RNA ligase (New England Biolabs #M2611A) according to the manufacturer’s instructions. For each sample, 10 µM of adenylated adapter was then ligated to the RNA template using a truncated KQ T4 RNA ligase 2 (New England Biolabs #M0373L) for 3 h at 25°C in the presence of 20% PEG8000 (New England Biolabs). Ligated RNA was gel-purified and size-fractionated on a 10% denaturing polyacrylamide gel to enrich for small RNA molecules in the size range of 50-400 nt using the low range ssRNA ladder as reference (New England Biolabs #N0364S). Gels were stained using 1x SYBR gold nucleic acid stain (ThermoFisher Scientific #S11494) in 1x TBE for 10 minutes and 3’adapter-ligated-RNA, with RNA sizes ∼50-400 nt, were excised on a blue light transilluminator. RNA was recovered from excised polyacrylamide gel pieces by crushing the gel fragments and soaking them into 0.3 M NaCl overnight at 4°C, followed by ethanol precipitation. For each sample, 5 µM of 5’ adapter with 4 terminal randomized nucleotides at the 3’ end was ligated using T4 RNA ligase (New England Biolabs #M0204S). Ligations were performed for 1 h at 25°C in the presence of 20% PEG8000. Thereafter, reverse transcription was performed for 1 h at 42°C using MarathonRT. Reactions were terminated by incubation with 0.25 M NaOH, for 3 min at 95°C, then neutralized with 0.25 M HCl. Next, cDNA was purified using MinElute Reaction Cleanup kit (Qiagen # 28006) and PCR amplified using Phusion high-fidelity PCR master mix (ThermoFisher Scientific # F531L) for 12 cycles of 98°C for 5 sec, 62°C for 10 sec and 72°C for 10 sec. Primers, randomized adapters, Illumina Multiplex and barcode primers were obtained from IDT and can be found in Scheepbouwer et al. 2023^58^. PCR products were equimolarly pooled for cluster generation without additional size-selection. Quality of sequence libraries was validated using 10% denaturing polyacrylamide gels, and Fragment Analyzer Systems (Agilent Technologies) was used to check size, purity and concentration. Sequencing of mid-range RNAseq was performed as paired-end reads for 150 cycles on NovaSeq 6000 at GenomeScan B.V. Leiden, Netherlands.

### Operational definition of pre-tRNAs and characterization

To enable high-confidence precursor boundary definition and downstream quantitative tRNA abundance measurements, two complementary analysis strategies were employed, consisting of a genome-based filtering approach for boundary characterization and a separate reference-based alignment strategy for quantification (see below).

Trimmed paired-end RNA-seq reads were aligned to the human genome (GRCh38) using STAR^92^ (v2.7.6a) with parameters configured to suppress spliced and chimeric alignments and to retain multimapping reads from repetitive loci (--outFilterMultimapNmax 100, --winAnchorMultimapNmax 100, --alignSJoverhangMin 200, --chimSegmentMin 10000). Alignments were processed using samtools, and subsequent analyses were performed using the Galaxy platform^93^ unless otherwise specified. Precursor-derived reads were operationally defined as reads overlapping annotated tRNA loci while extending beyond mature tRNA boundaries. Reads confined to the mature tRNA body are frequently ambiguous due to sequence redundancy, whereas extension into flanking regions increases locus specificity and enables identification of precursor-derived fragments. Genome-aligned BAM files were subjected to a two-step filtering strategy. First, alignments were filtered using deepTools^94^ alignmentSieve (v3.5.4), restricting fragment length to ≤400 nt and retaining first-mate alignments from properly paired reads (SAM flag 66). Because tRNA and pre-tRNA species are short relative to the 150 bp paired-end read length, informative precursor-derived fragments are expected to be recovered as overlapping proper pairs, whereas singleton or improperly paired reads are more likely to reflect incomplete or low-confidence fragment recovery and were therefore excluded from boundary definition analyses. To exclude mature tRNA-derived reads, a custom blacklist was applied consisting of annotated tRNA loci extended by 1 nt upstream and 3 nt downstream, accounting for minor imprecision at 5′ processing sites and post-transcriptional CCA addition at the 3′ end. Second, alignments were filtered using samtools^95^ (v1.8) to retain only reads overlapping annotated tRNA loci. Together, these steps ensure retention of reads anchored within tRNA genes while extending beyond annotated mature boundaries, while excluding fragments confined to mature tRNA bodies or mapping exclusively to flanking regions. Filtered alignments were used to generate base-resolution coverage tracks using deepTools^94^ bamCoverage (v3.5.4) with 1 nt binning and CPM normalization. Gene body–scaled coverage profiles were generated using deepTools^94^ computeMatrix (v3.5.4) in scale-regions mode, with annotated tRNA loci scaled to 72 nt (mature tRNA body) and including 40 nt upstream and 120 nt downstream flanking regions. Regions were sorted by mean signal intensity, and missing values were set to zero. Heatmaps and average profiles were generated using deepTools^94^ plotHeatmap (v3.5.4). Because the filtering strategy enriches for reads extending beyond mature boundaries, coverage profiles predominantly reflect precursor-derived fragments and processing intermediates, enabling resolution of processing features at tRNA boundaries. Coverage matrices were exported for downstream quantitative analysis and operational definition of pre-tRNA boundaries. Quantitative tRNA abundance measurements were performed separately using a permissive reference-based alignment strategy as described below.

#### Boundary definition and processing-state quantification

Genome-wide coverage profiles were highly consistent across biological replicates. Inspection of individual loci revealed sharp and reproducible transitions at the 5′ boundary of the coverage profile and at the transition into the mature tRNA region (leader processing), as well as signal decay downstream of the mature region (trailer). Transitions associated with 3′ trailer processing were less abrupt, consistent with the expected order of processing. For quantitative boundary definition, coverage values for each locus and replicate were normalized to the mean signal across the scaled mature tRNA body (72 nt). Positions were defined relative to the mature tRNA boundaries, with position −1 corresponding to the last nucleotide upstream of the mature region and +1 the first nucleotide within the mature tRNA. Normalized profiles were aggregated across loci to generate genome-wide positional coverage profiles for each time point. Genes with zero counts in any sample were excluded. The 5′ boundary was defined as the upstream position at which normalized coverage decreased by ≥90% relative to position −1. Leader processing fractions were quantified at position −1 relative to the mature boundary. To define transcription termination, coverage decay downstream of position +73 (first nucleotide of the trailer region) was evaluated. Because terminators may display variable levels of transcription read-through^63^, candidate termination positions were defined by ≥50% (T50), ≥90% (T90), and ≥97% (T97) reductions in normalized coverage relative to +73. Primary termination sites (T1) were assigned at T90 when T50 and T90 occurred within 4 bp of each other, and otherwise at T50. Trailer processing fractions were quantified at position +73 relative to the mature boundary (+72). Because terminators occur at defined sequence motifs, T1 positions were further evaluated based on local T content. Canonical terminators were defined as sites where T1 corresponds to the last nucleotide of a stretch of ≥4 consecutive Ts. In most loci, T1 matched this position directly, with assignments typically within ±1 bp (90%). Sites lacking such stretches were classified based on local T content: positions containing ≥4 non-contiguous Ts within a 6 bp window were defined as non-canonical terminators (6%), while sites lacking these features, including absence of ≥3 consecutive Ts within an 8 bp window, were classified as undetermined sites (4%). These sites were associated with extended trailer regions (median 55 nt versus 13 nt for canonical terminators). To quantify read-through transcription while minimizing sensitivity to positional imprecision in T1 assignment, signal was measured 4 bp downstream of T1. This offset captures residual transcription beyond termination sites while reducing contributions from gradual signal decay across termination regions.

### Quantitative tRNA expression analysis and precursor-to-mature ratio measurements

A custom tRNA reference sequence library was constructed by integrating mature, precursor, and mitochondrial tRNA sequences. Mature nuclear tRNAs were obtained from GtRNAdb^96^ and converted to DNA alphabet; a canonical 3′ CCA tail was appended to reflect post-transcriptional maturation. Identical sequences were collapsed based on sequence identity while retaining all associated gene annotations, and the number of genes per collapsed sequence was encoded in the FASTA header. Precursor tRNA sequences were derived from hg38 genomic loci using the operational definition described above and similarly collapsed without CCA extension. Mitochondrial tRNAs were obtained from RNAcentral (HGNC-linked entries) and extended with a 3′ CCA tail. The final reference library consisted of concatenated mature nuclear, precursor, and mitochondrial tRNA sequences. To enable deterministic resolution of multimapping reads, the reference was ordered with mature nuclear tRNAs first, followed by precursor and mitochondrial tRNAs. Reads lacking both precursor-specific sequences and a 3′ CCA tail can align equivalently to mature and precursor reference sequences, particularly for intronless loci. Because the CCA tail can be removed and re-added post-transcriptionally, absence of CCA does not uniquely indicate precursor state. This reference ordering was designed to preferentially assign such reads to mature tRNAs. Considering the short half-life of precursor tRNAs relative to mature tRNAs, this assignment is also expected to reflect the predominant origin of CCA-less molecules.

Adapter sequences were removed using cutadapt^97^ (v5.2) with custom adapter definitions containing degenerate positions and a minimum overlap of 9 nt (including at least 5 non-degenerate positions). Quality trimming was performed using --nextseq-trim=20 to account for poly(G) artifacts. Trimmed paired-end reads were aligned to the custom reference using BBMap^64,65^ (v39.08) in local alignment mode (local=t), allowing short insertions and deletions up to 5 nt (maxindel=5). A permissive alignment identity threshold was used to retain reads containing mismatches arising from reverse transcription across modified bases. Reads with multiple equally scoring alignments were assigned to the first best site (ambiguous=best) under deterministic mode (deterministic=t), ensuring reproducible assignment consistent with the predefined reference hierarchy. Read counts per reference sequence were extracted using samtools^95^ idxstats (Galaxy tool v2.0.8) and normalized to counts per million (CPM). All downstream analyses were performed on CPM-transformed values. Across samples, reads with multiple equally best alignments represented a small fraction of total reads (mean ± SD: 4.7 ± 1.3%), indicating that only a limited subset of reads was subject to deterministic multimapping resolution. Absence of major bias introduced by permissive alignment parameters and deterministic multimapping resolution, was confirmed via comparisons of the results to ALL-tRNAseq pipeline^58^ analyses of mature cytoplasmic and mitochondrial tRNAs using Bowtie followed by Smith-Waterman refinement.

Global tRNA composition was assessed by aggregating reads assigned to cytoplasmic (mature and precursor) and mitochondrial tRNAs and computing their relative fractions within each sample, followed by averaging across replicates for each time point. tRNAs with low expression (CPM < 2 across time points) were excluded prior to log transformation. Principal component analysis (PCA) and pairwise Pearson correlation coefficients were calculated on log_2_-transformed CPM values across all tRNA species to assess global expression patterns and evaluate reproducibility between biological replicates prior to averaging (see **Figure 5C**). PCA was performed using the Galaxy tool ggplot2_pca (v3.5.1), which implements PCA via ggfortify^98^ using default settings. To distinguish coordinated shifts in relative abundance from gene-specific changes, distribution-level differences across time points were assessed using the Friedman test applied to matched tRNA expression profiles. In parallel, preservation of relative expression was evaluated by pairwise Pearson correlation analysis within tRNA classes (mature cytoplasmic, mitochondrial, and precursor tRNAs) using replicate-averaged values per time point.

Precursor-to-mature ratio (*R*) analyses were restricted to uniquely assignable precursor–mature pairs. Pairing was established by matching precursor gene annotations to the gene annotations retained for collapsed mature tRNA sequences. tRNAs were classified as unique when a single precursor could be assigned unambiguously to a single mature tRNA species. Cases in which multiple precursor genes mapped to the same mature sequence, or in which precursor loci could not be distinguished due to sequence identity, were excluded. Pairs with low precursor expression (CPM ≤ 10 in replicate-averaged values at any time point) were also excluded to avoid instability in ratio estimates arising from low-count noise and log transformation. For each unique pair, *R* was calculated within each replicate, averaged across replicates at each time point, and expressed as log_2_-transformed ratios. For comparison across time points, ratios were baseline-normalized to the differentiated state (0 h) as log_2_(*R*/*R_0h_*). The fraction of tRNAs exhibiting reduced ratios at 48 h relative to 0 h was quantified as the proportion of pairs with log_2_(*R_48h_*/*R_0h_*) < 0 (equivalently, *R_48h_* < *R_0h_*).

### Dynamic and steady-state constraints on precursor-to-mature tRNA ratios *R*

The precursor-to-mature ratio (*R* = *P* / *M*) reflects the balance between the generation of mature tRNAs through precursor processing and their loss through degradation and dilution associated with cell growth. The dynamics of the mature tRNA pool are described by *dM*/*dt* = *k_1_·P* − (*k_degM_ + g*)·*M*, where *k_1_*is the effective rate constant for precursor-to-mature processing, *k_degM_*is the effective degradation rate constant of mature tRNAs, and *g* is the growth rate accounting for dilution during cell proliferation. Dividing by M gives:

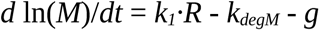

This relationship holds independently of whether the system is at steady state and shows that changes in mature tRNA abundance depend on the balance between precursor processing, mature tRNA degradation, and growth-associated dilution.

At steady state, the rate at which mature tRNAs are produced must match the rate at which they are lost. This gives: *k_1_*·*P* = (*k_degM_* + *g*)·*M*, where *k_1_* is the effective rate constant for precursor-to-mature processing, *k_degM_* is the effective degradation rate constant of mature tRNAs, and *g* is the cell growth rate, which accounts for dilution of RNA species due to cell division. Rearranging this expression yields: *R* = *P*/*M* = (*k_degM_* + *g*) / *k_1_*. In non-proliferating differentiated cells (*g* = 0), this reduces to *R_diff_* = *k_degM_* / *k_1_*. Under steady-state proliferation (*g* > 0), the ratio becomes *R_prolif_* = (*k_degM_* + *g*) / *k_1_*. Thus, if processing and decay remain unchanged, proliferation is expected to increase R due to dilution of the mature tRNA pool such that *R_diff_* < *R_prolif_*.

These steady-state relationships are independent of transcriptional activity and precursor degradation. However, they do not necessarily describe the transient dynamics during re-entry into proliferation, particularly for long-lived mature tRNAs. Changes in transcription may temporarily alter *R* even when processing and decay rates remain constant. We therefore used numerical simulations to evaluate whether transcriptional regulation alone could reproduce the experimentally observed reductions in *R* under non-steady-state conditions.

### Mathematical modeling and simulation of *R* dynamics

To assess whether changes in *R* can be explained by transcriptional regulation alone, we employed a two-compartment kinetic model describing precursor and mature tRNA dynamics. The model incorporates transcription, precursor-to-mature processing, RNA decay, and dilution due to cell growth. Under constant post-transcriptional parameters (*k_1_* and *k_degM_*), the model is consistent with the steady-state constraint derived above.

To evaluate whether transient, non-steady-state dynamics could nevertheless produce reductions in *R*, we performed numerical simulations across a broad range of time-dependent transcriptional programs, while keeping all post-transcriptional parameters constant. Simulations were initialized at steady state and evaluated at time points matching experimental sampling (0, 12, and 48 h), and changes in *R* were quantified relative to the initial condition. In parallel, scenarios incorporating time-dependent changes in precursor processing or mature RNA decay were simulated to assess the contribution of post-transcriptional regulation. Comparison of these regimes was used to determine whether transcription alone can account for the observed decrease in *R* or whether modulation of post-transcriptional parameters is required. A full mathematical formulation and detailed description of the simulation framework are provided in **Supplementary Note S1**.

### Gene set enrichment analyses (GSEA)

Total and polysomal RNA microarray data from differentiated and proliferative HepaRG cells (ArrayExpress E-MEXP-1082)^44^ were preprocessed using the Robust Multi-chip Analysis (RMA) algorithm for background adjustment, quantile normalization and summarization with Transcriptome Analysis Console (TAC, v4.0.3.14; ThermoFisher Scientific). TAC differential expression output tables were used for gene set enrichment analysis. RNA sequencing data from human induced hepatocyte (hiHep)-derived organoids overexpressing GFP or MYC (Gene Expression Omnibus GSE115853)^68^ were analyzed using GEO2R (NCBI), which applies the limma framework^99^ (v3.54.0) to processed GEO Series matrix data. Filtering thresholds were relaxed (FDR = 0.9999, |log_2_FC| = 0) to retain the full gene set for downstream preranked GSEA.

For HepaRG data, TAC differential expression tables were processed to generate preranked gene lists for WEB-based GEne SeT AnaLysis Toolkit (WebGestalt)^100^. Cross-hybridizing probe sets (_x_at), entries without gene symbols, and probes annotated to multiple genes were removed. Probe sets mapping to the same gene were collapsed by retaining the probe set with the lowest FDR-adjusted P value. A preranking score was calculated from the raw P value as -log _10_(P) × sign(log_2_ fold change), using the raw P value and direction of expression change. Genes were ranked accordingly for preranked GSEA using WebGestalt^100^ with the Reactome database. Differential expression results from MYC-overexpressing organoids were similarly used for Reactome GSEA.

For the initial HepaRG analysis, significantly enriched Reactome terms were considered at FDR < 0.05. To visualize enrichment patterns across RNA-related processes, enriched terms were further selected by keyword matching against RNA-associated annotations, including terms related to RNA, mRNA, rRNA, tRNA, RNA degradation, and RNA polymerase III. Filtered terms were visualized using normalized enrichment scores (NES) and FDR-based summary plots, with tRNA- and RNA polymerase III-related pathways displayed separately. For the MYC-overexpression dataset, the 20 most strongly enriched Reactome pathways were selected based on NES.

### Codon usage analysis

MANE Select^101^ transcript coding sequences (CDS) were retrieved from Ensembl (via BioMart) to obtain a single representative transcript per gene. Codon counts were computed from CDS sequences parsed in-frame, excluding the terminal stop codon. Internal TGA codons were retained to accommodate selenocysteine-encoding transcripts, while TAA and TAG stop codons were excluded from codon usage comparisons. Codon counts were aggregated at the gene level to generate a codon count matrix.

Normalized total and polysomal RNA expression values from the HepaRG TAC analysis were used to estimate transcript abundance. Cross-hybridizing probes (_x_at) and probes without gene symbols were excluded, and probe sets mapping to the same gene were collapsed by taking the median of their log_2_-transformed expression values. These values were subsequently converted to linear scale for downstream analyses, providing a robust estimate of gene-level expression independent of probe-level statistical variation.

Genes present in both the codon count matrix and expression dataset were retained for analysis. Codon demand for each condition was calculated as the expression-weighted sum of codon counts across all genes, such that codon usage reflects the contribution of each transcript scaled by its abundance. Resulting codon counts were normalized to relative codon frequencies within each condition. Codon enrichment between conditions was quantified as log2 fold change in relative codon usage. For comparisons between polysomal and total RNA fractions, enrichment was defined as log_2_(Poly/Total) for both proliferative and differentiated states.

Statistical significance of codon enrichment was assessed using an expression-stratified permutation approach. Genes were stratified into expression deciles based on their average expression across proliferative total and polysomal RNA fractions. Expression values from each fraction were independently permuted among genes within each decile to preserve the overall expression distribution. For each of 10,000 permutations, codon demand and corresponding log2 fold changes were recomputed. Empirical two-sided P values were calculated as the proportion of permutations yielding an absolute log2 fold change at least as large as the observed value, with a correction to avoid zero P values. P values were adjusted for multiple testing using the Benjamini–Hochberg procedure.

### PerturbAtlas analysis

To identify potential regulators of the tRNA processing program, the 47 leading-edge genes associated with the Reactome “tRNA processing” pathway (R-HSA-72306) in proliferative HepaRG cells were retrieved from WebGestalt. Gene-level log_2_ fold changes (proliferative versus differentiated) were obtained from TAC total RNA differential expression data. Cross-hybridizing probes (_x_at), probes without gene symbols, and probes mapping to multiple genes were excluded. For genes represented by multiple probes, the probe with the largest absolute log_2_ fold change among those with FDR < 0.05 was selected, or the probe with the lowest FDR if none met this threshold. This approach prioritized statistically supported expression changes rather than representative transcript abundance. The resulting 47 genes were mapped to Ensembl identifiers and submitted with their corresponding log_2_ fold changes to the Gene Rank analysis module of PerturbAtlas^83^.

PerturbAtlas enrichment results were downloaded and combined for downstream analysis. Only significantly enriched perturbations (FDR < 0.05) showing directionality consistent with positive regulation of the tRNA processing program were retained: gain-of-function perturbations (overexpression or activating mutations) with NES > 1 and loss-of-function perturbations (knockdown or knockout) with NES < -1. Perturbation targets were matched across gain- and loss-of-function datasets, and only targets represented in both groups were retained. The resulting 48 perturbation records, corresponding to 15 targets, were visualized using NES and -log_10_(adjusted p-value).

### Comparison of HepaRG and MYC-overexpression transcriptomic datasets

To assess whether the tRNA biogenesis program associated with proliferation in HepaRG cells is recapitulated by MYC overexpression, Reactome GSEA results from both datasets were compared. WebGestalt results were exported without significance filtering or redundancy reduction to retain enrichment statistics and leading-edge gene sets for all Reactome terms. For comparison of tRNA- and RNA polymerase III-related pathways, terms significantly enriched (FDR < 0.05) in at least one dataset were selected. Enrichment scores from both datasets were compared.

For gene-level comparisons, leading-edge genes associated with the Reactome tRNA processing pathway were retrieved from WebGestalt output and merged with independently generated log_2_ fold-change tables for each dataset. HepaRG gene-level fold changes were obtained from TAC total RNA data using the probe-selection procedure described above for PerturbAtlas, applied to all genes. MYC-overexpression fold changes were obtained from GEO2R differential expression results. A heatmap was generated using the union of leading-edge genes identified in either dataset, retaining only genes with available expression measurements in both datasets. Log_2_ fold changes were standardized within each dataset using z-scores calculated from the global distribution of gene-level log_2_ fold changes across all genes, allowing comparison between datasets with different dynamic ranges while preserving the direction of expression changes. Genes were annotated according to whether they belonged to the leading edge in HepaRG, MYC-overexpressing organoids, or both.

### Data visualization and statistical analysis

Simulations were carried out in Python (v3.13.9) using standard scientific libraries (NumPy, SciPy, Matplotlib) within the Spyder environment (6.1.4). Sequencing data were processed and analyzed on the Galaxy platform^93^ and visualized with IGV^102^ (2.19.7). Microarray data were analyzed with Transcriptome Analysis Console (TAC, v4.0.3.14). Gels and blots were quantified with GelAnalyzer^103^ (23.1.1) and FIJI ImageJ^104^ (1.54r). Statistical analyses were conducted in JASP^105^ (0.96.0) and growth or decay curve fitting was performed with LabPlot^106^ (2.12.1) and Gnumeric^107^ (1.12.60). For WebGestalt^100^, PerturbAtlas^83^, and Enrichr^84^ analyses, statistics reported by the respective tools were used.

## Data and code availability

RNA-seq data have been deposited at Gene Expression Omnibus (GEO) and will be publicly available as of the date of publication. This paper also analyzes existing, publicly available data, accessible at GSE115853 (GEO) and E-MEXP-1082 (ArrayExpress). Any additional information required to reanalyze the data reported in this paper is available from the lead contact upon request.

## Acknowledgments

We are particularly grateful to Robert G. Roeder for his generous support, and contributions to this project. All experiments investigating RNA polymerase III isoforms were performed by A.G. in his laboratory at The Rockefeller University. We also gratefully acknowledge the antibodies, cell lines, and other reagents provided by his laboratory that were instrumental to this work. We greatly appreciate his continued encouragement and support throughout the development of this study.

We thank Ueli Schibler, Evelina Tutucci, Keiichi Ito, Zeynep Veryeri and Gerben Vader for their comments on the manuscript. This work was supported by a NWO Talent Programme Vidi grant [VI.Vidi.193.107], an Amsterdam UMC Starter Grant [4.04.127_SdJ.kb], a Stichting Cancer Center Amsterdam [CCA2021-5-26] and a ZonMW Open competition grant [09120232310068]. A.G. was also supported by a Swiss National Science Foundation Early Mobility Fellowship [P2GEP3_151952] and by a Human Frontier Science Program Long-Term Fellowship [LT001083/2014] for experiments performed at the Rockefeller University.

## Author Contributions

M.G. and D.S. designed and performed experiments and analyzed data. C.S. established and performed the tRNA sequencing experiments. B.G.O. and B.S. contributed to experimental validation and data analysis. C.G.M and M.H. performed initial sequencing data analysis. D.M.P and M.H supervised part of the work performed. A.G. designed the project, acquired funding, supervised and performed experiments, analyzed, visualized and interpreted the data, developed and implemented the final bioinformatics pipeline and the theoretical analysis framework and wrote the manuscript. All authors read the manuscript.

## Supplementary Information

## 1 Supplementary Figure legends

**Figure S1.**
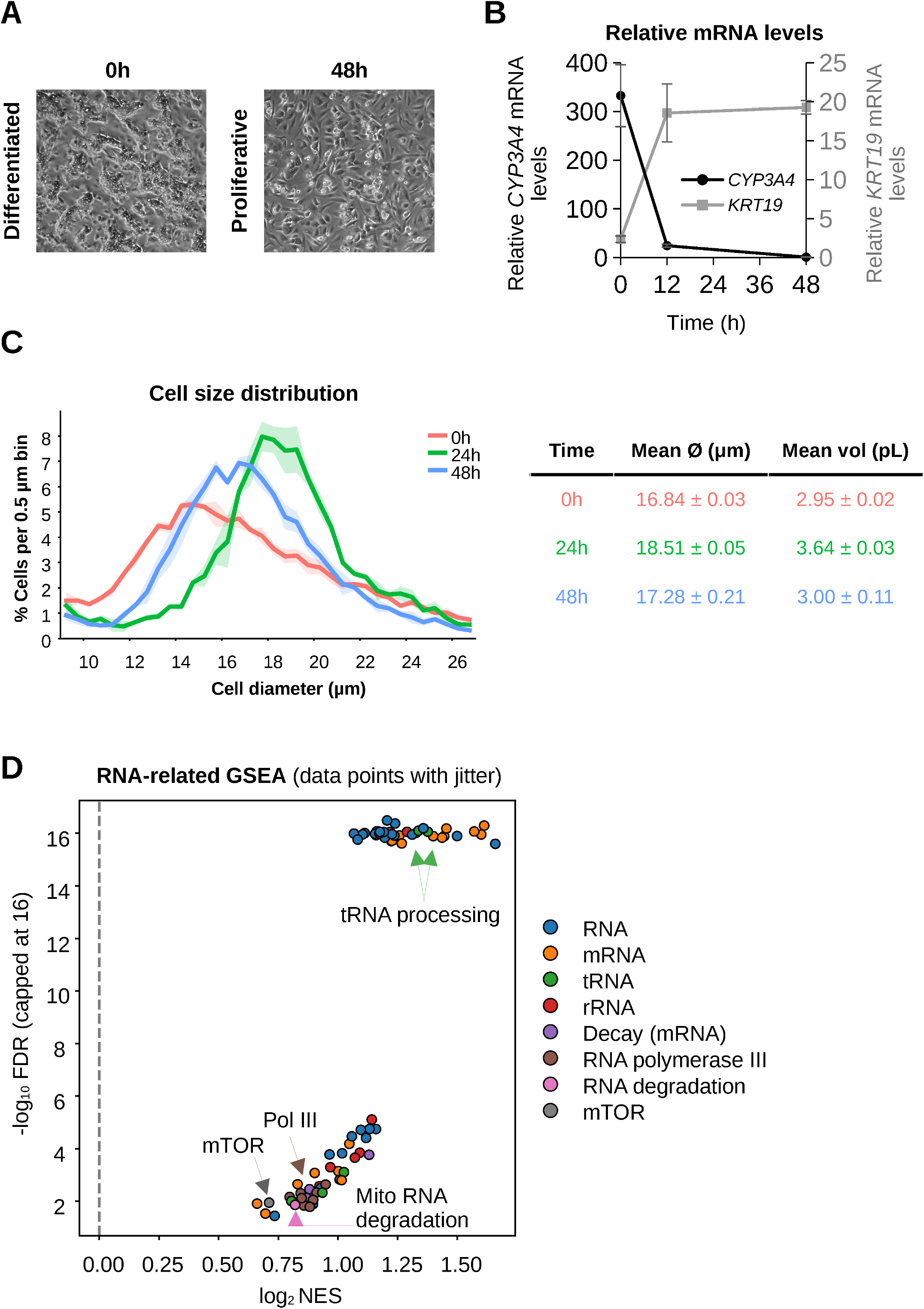
Related to Figure 1. **A)** Representative phase-contrast images of differentiated (0 h) and proliferative (48 h) HepaRG cells following replating at low density. **B)** RT-qPCR analysis of *CYP3A4* and *KRT19* mRNA levels relative to *TBP* at indicated time points (mean *±* SD, *n* = 3). **C)** Cell size distributions of differentiated HepaRG cells (0 h) and cells at 24h and 48h following replating, measured using a CASY cell counter. Distributions represent the percentage of cells per 0.5 *µ*m diameter bin, normalized to the total number of cells within the 9-27 *µ*m diameter range. The accompanying table shows mean cell diameter and estimated cell volume (mean *±* SD) from three independent biological experiments, each measured in technical duplicate. Technical measurements were combined within each biological replicate. **D)** Gene set enrichment analysis (GSEA) of RNA-related pathways. Only significantly enriched pathways (FDR < 0.05) are shown. Each point represents a gene set plotted by log_2_-transformed normalized enrichment score (NES) versus *−* log_10_(FDR), capped at 16 for visualization. Selected pathways are indicated. Data points are jittered to reduce visual overlap.

**Figure S2.**
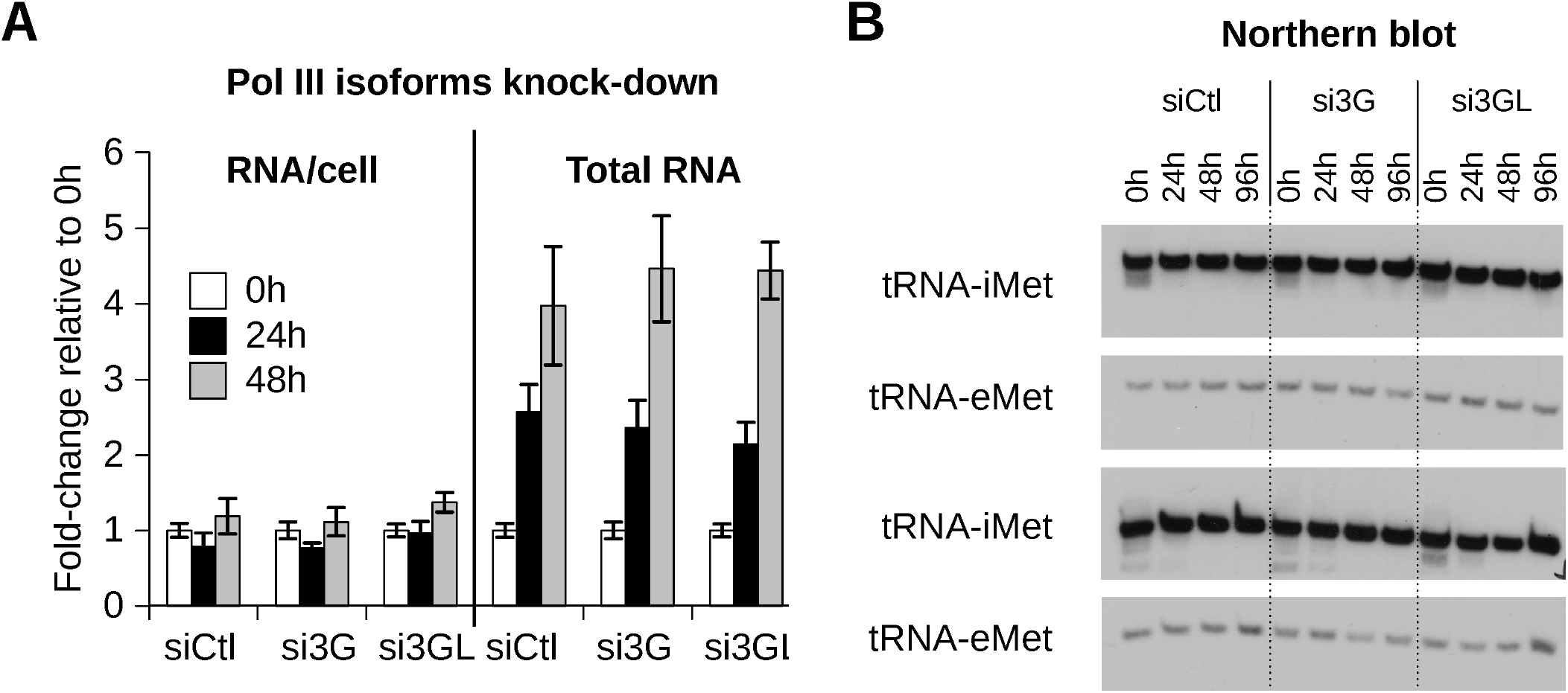
Related to Figure 2. **A)** RNA content per cell (left) and total RNA per culture (right) at indicated time points following siRNA-mediated depletion of *POLR3G* (si3G), *POLR3GL* (si3GL), or control (siCtl). Values are shown as fold-change relative to 0 h. Data represent mean *±* SD (*n* = 3). **B)** Northern blot analysis of cytoplasmic tRNAs (tRNA-iMet and tRNA-eMet) across time (0 h, 24 h, 48 h, 96 h) following isoform depletion. Data correspond to quantification shown in Figure 2E.

**Figure S3.**
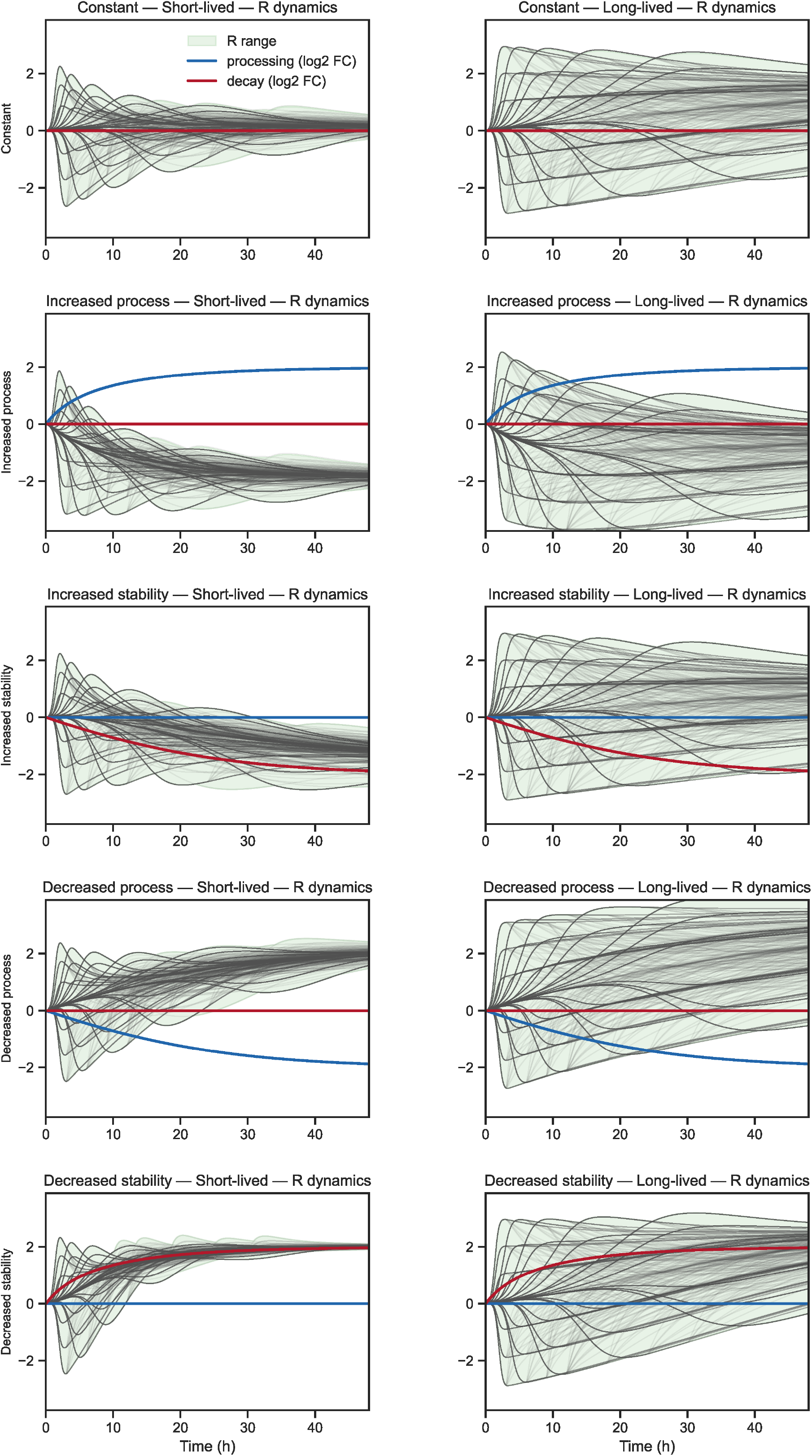
Related to Figure 3. Simulated trajectories of log_2_(*R/R*_0_*_h_*) over time for short-lived (left) and long-lived (right) mature tRNAs under distinct parameter regimes. Conditions shown are constant parameters, increased processing (increased *k*_1_), increased stability (reduced *k*_degM_), decreased processing, and decreased stability. Individual trajectories are shown in gray. Colored lines indicate the time-dependent log_2_fold changes in the processing rate (*k*_1_, blue) and mature tRNA degradation rate (*k*_degM_, red), overlaid on the simulated precursor-to-mature ratio trajectories.

**Figure S4.**
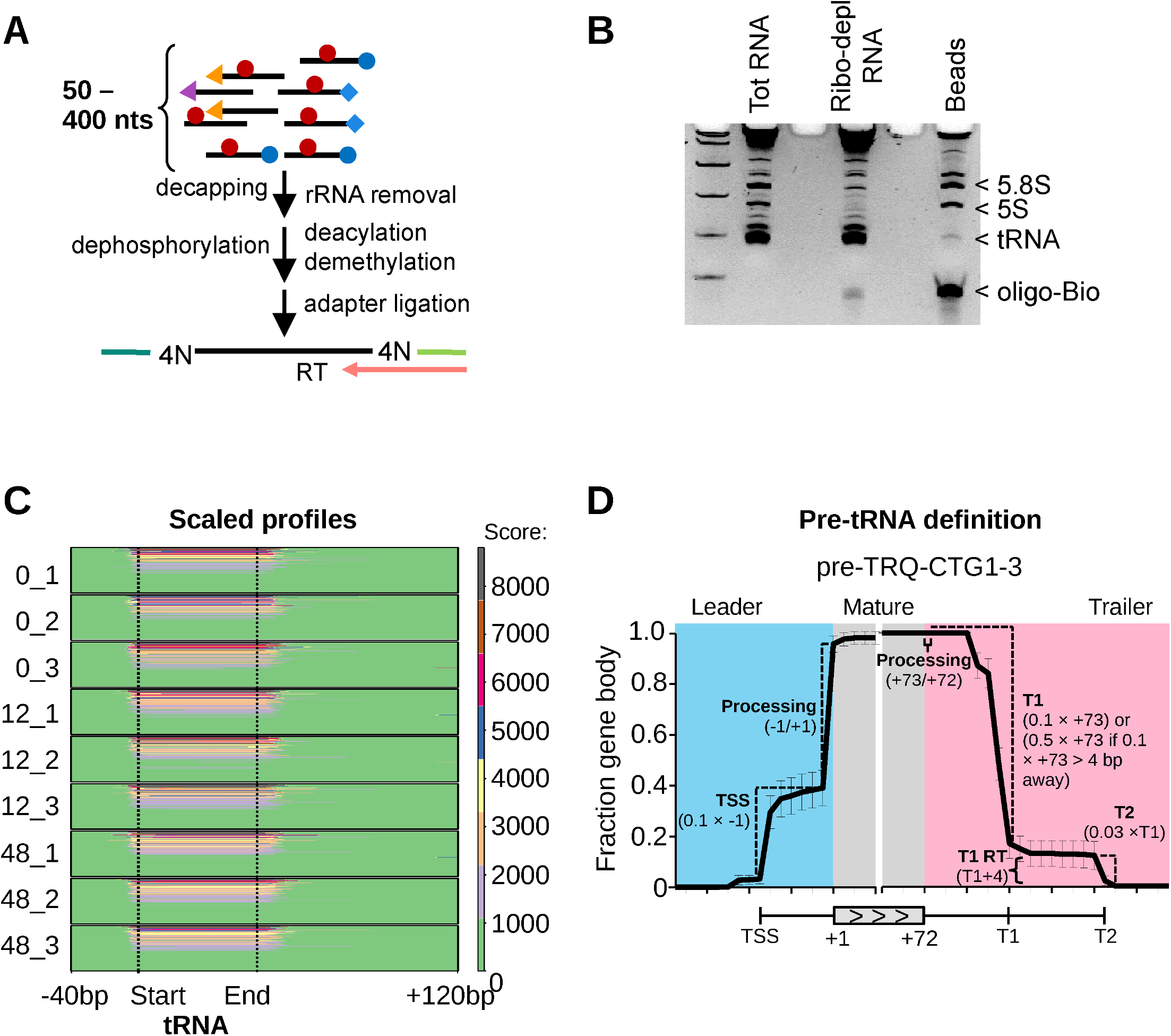
Related to Figure 4 and 6. **A)** Schematic of tRNA sequencing library preparation, including RNA pre-treatments, adapter ligation, reverse transcription. Colored symbols schematically represent RNA terminal structures (5 caps, triphosphates, and 3 aminoacylation) and internal modifications (methylation) targeted by the indicated pretreatments. **B)** Representative denaturing gel showing depletion of 5S and 5.8S rRNA by hybridization followed by pull-down. Total RNA, rRNA-depleted RNA, and bead-bound fractions are shown. **C)** Scaled coverage profiles across all individual tRNA genes and replicates used for boundary detection. Coverage is normalized to the mature tRNA body and aligned relative to annotated start and end positions. **D)** Example locus illustrating the operational definition of pre-tRNA features, including transcription start site (TSS), leader, mature tRNA body, trailer, and transcription termination sites (T1 and T2). Processing and transcriptional boundaries are inferred from normalized coverage profiles and used for genome-wide annotation.

**Figure S5.**
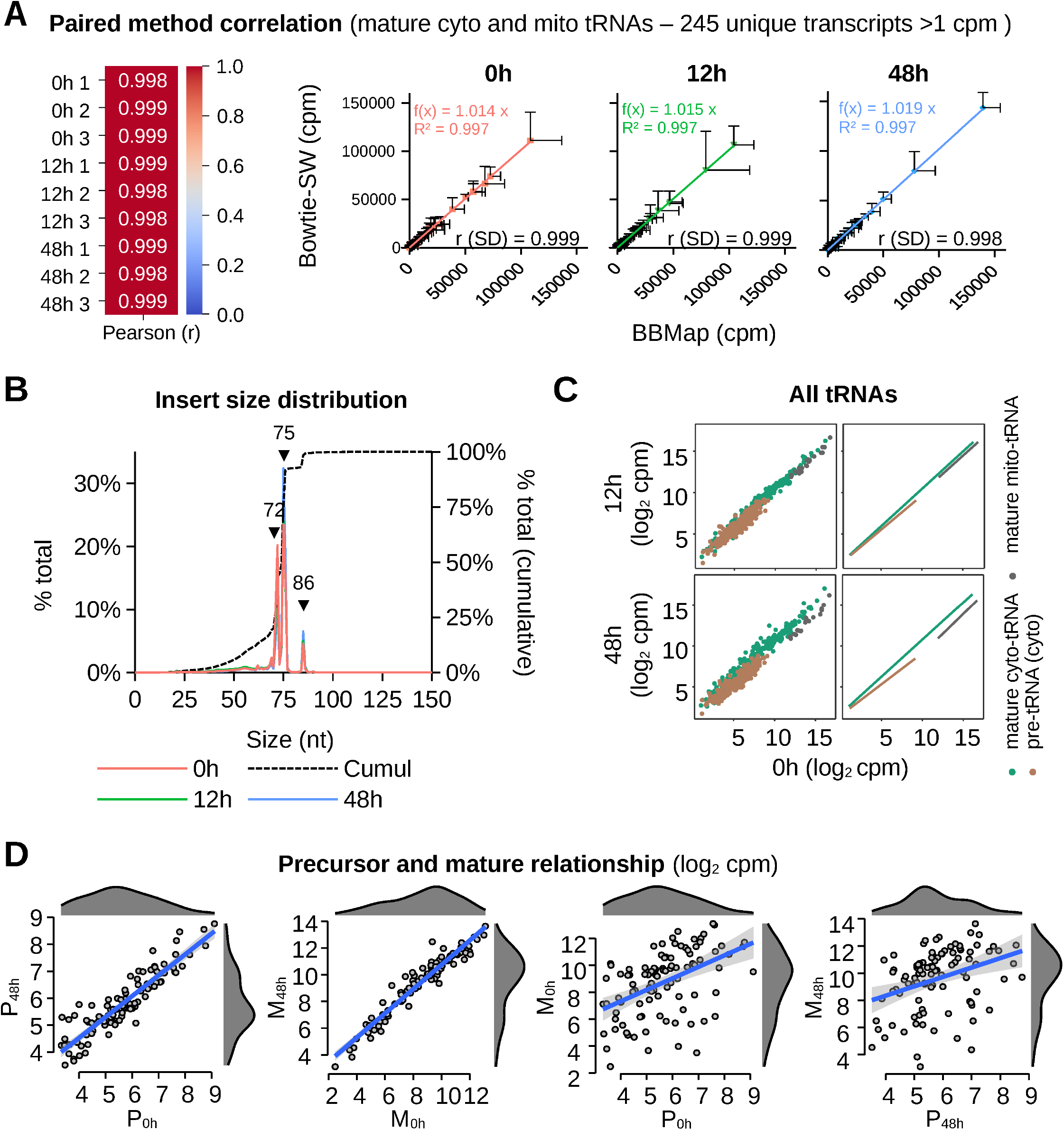
Related to Figure 5. **A)** Pairwise correlation of tRNA expression estimates obtained from independent mapping strategies (BBMap and Bowtie) for uniquely assignable (i.e., identical sequences collapsed) mature cytoplasmic and mitochondrial tRNAs (>1 cpm). Pearson correlation coefficients are shown for all replicates and time points. **B)** Insert size distribution up to 150 nts of the aligned paired-end reads at indicated time points. Cumulative fraction is shown on the right y-axis. **C)** Pairwise comparison of expression levels (log_2_ cpm) across time points for cytoplasmic mature tRNAs (green), precursor tRNAs (brown), and mitochondrial tRNAs (grey). **D)** Relationship between unique precursor and mature tRNA expression (log_2_ cpm; average of three replicates) across time points. Scatter plots show pairwise comparisons between precursor and mature levels within and across time points.

**Figure S6.**
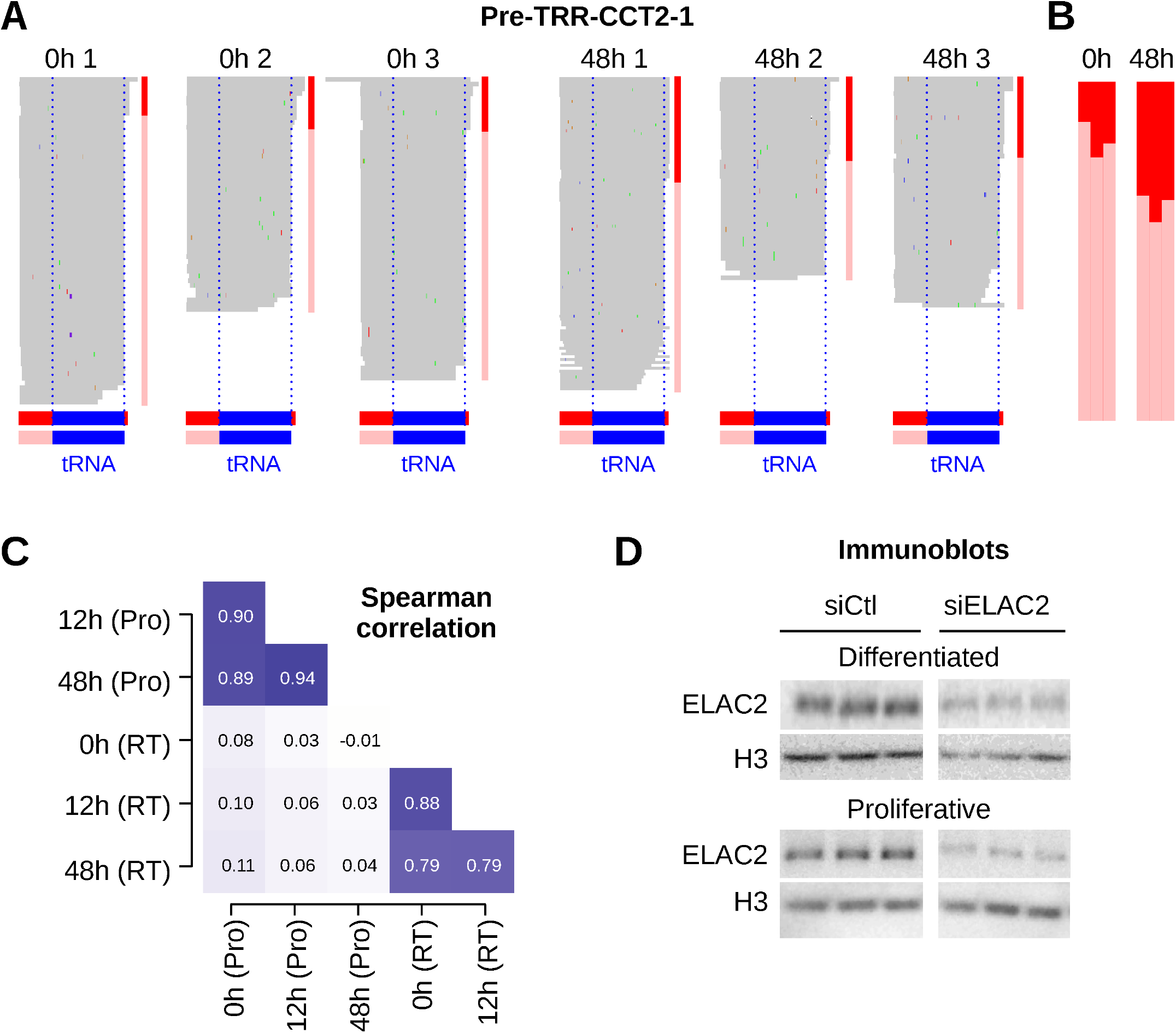
Related to Figure 6. **A)** IGV genome browser views of an example tRNA locus (pre-TRR-CCT2-1) showing filtered read alignments across replicates at 0 h and 48 h. Reads are sorted by length and used to distinguish leader-containing versus leader-processed species. This alignment-based representation is independent of the coverage-based quantification. This locus was selected for dynamic range; similar changes are observed across loci with detectable baseline processing (Figure 6). **B)** Fraction of precursor reads retaining an unprocessed 5 leader at the indicated locus, relative to all precursor reads. **C)** Spearman correlation between the fraction of leader-unprocessed precursors (Pro) and T1 read-through (RT) levels across genes with detectable baseline processing (⩾5% at 0 h) and reproducible measurements (SD < 0.2 across replicates at all analyzed time points for both Pro and RT; 98 genes), at indicated time points. Related to Figure 6B**–C**. **D)** Immunoblot analysis of ELAC2 in differentiated and proliferative HepaRG cells following siRNA-mediated depletion of *ELAC2* (siELAC2) or control (siCtl). Histone H3 is shown as loading control.

**Figure S7.**
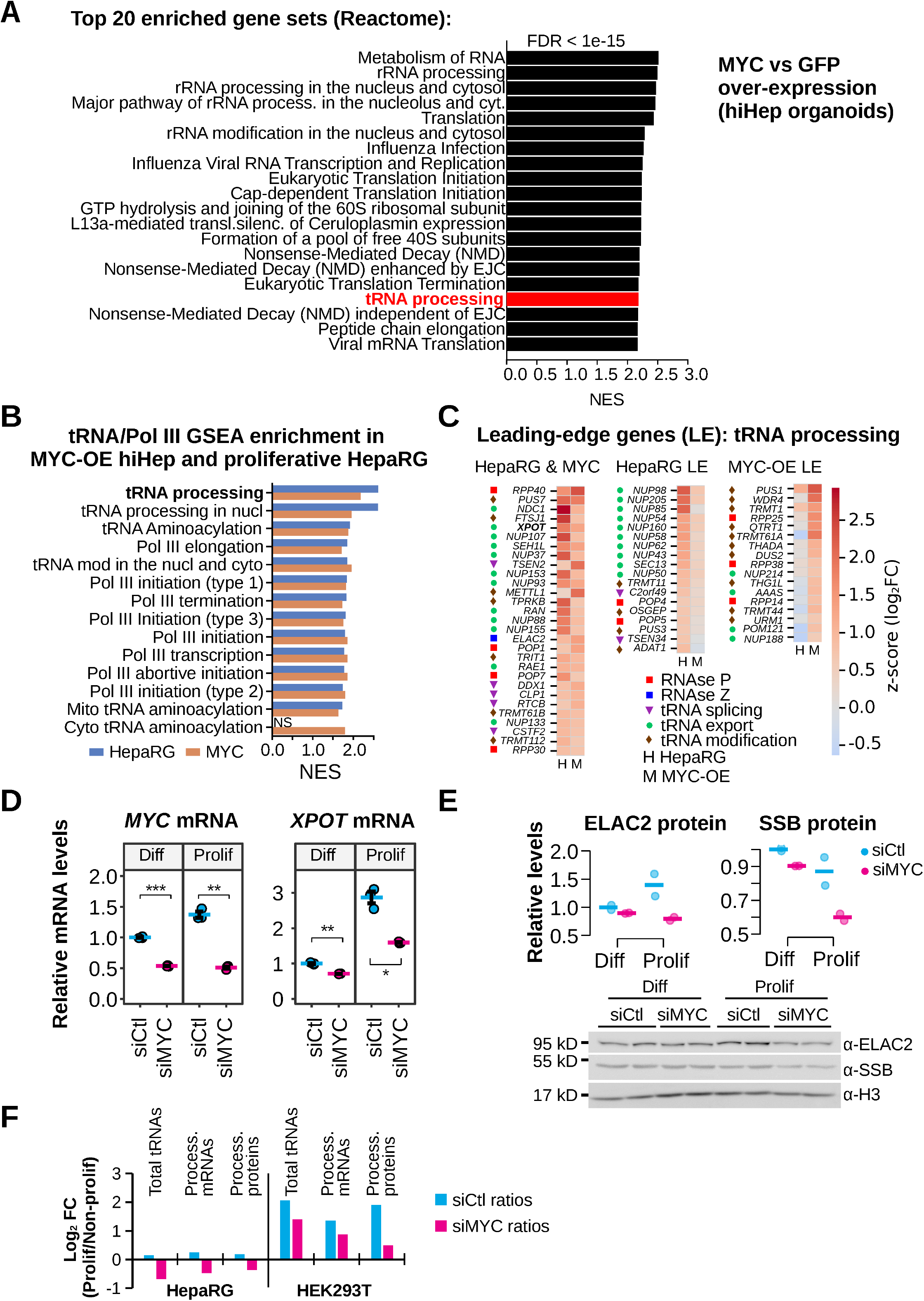
Related to Figure 7. **A)** Gene set enrichment analysis (GSEA) of Reactome pathways in MYC-overexpressing hiHep organoids. The top 20 enriched pathways are ranked by normalized enrichment score (NES); all displayed terms have FDR < 1*×*10^−15^. **B)** Gene set enrichment analysis (GSEA) of tRNA- and Pol III-related pathways in MYC-overexpressing hiHep organoids and proliferative HepaRG cells. NS, non-significant (FDR *≥* 0.05). **C)** Leading-edge genes (LE) contributing to enrichment of the tRNA processing pathway in HepaRG (H), MYC-overexpressing hiHep organoids (M), and their overlap. Heatmap shows expression changes (Z-scores of log_2_ fold-change) across conditions. Functional categories (RNase P, RNase Z, tRNA splicing, export, and modification) are indicated. **D)** RT-qPCR analysis of *MYC* and *XPOT* mRNA levels relative to *TBP* (*n* = 3) in differentiated (Diff) and proliferative (Prolif) HepaRG cells following control (siCtl) or *MYC* knockdown (siMYC). Values are normalized to siCtl Diff (control differentiated). Data represent mean *±* SD. Statistical significance was assessed using Welch’s t-test (*p < 0.05; **p< 0.005; ***p<0.0005). **E)** Immunoblot analysis of ELAC2 and SSB protein abundance in differentiated and proliferative HepaRG cells following control or *MYC* knockdown, with corresponding quantification (*n* = 2). Proteins were recovered from the organic phase of the same TRIzol lysates used for RNA quantification (Figure 7F and **7H**). Histone H3 was used as a proxy for DNA/cell-number loading and to normalize ELAC2 and SSB protein abundance. Values are expressed relative to siCtl differentiated cells. **F)** Summary of proliferation-associated changes in total mature tRNA abundance, processing-factor mRNAs, and selected processing proteins in HepaRG and HEK293T cells following control or *MYC* knockdown. Bars represent the log_2_-transformed arithmetic mean fold-change between proliferative and non-proliferative conditions for each molecular category, calculated separately for siCtl and siMYC. Processing-factor mRNAs comprise *SSB*, *RPP40*, *ELAC2*, *TSEN2*, and *XPOT*, and processing proteins comprise ELAC2 and SSB. Values were derived from matched per-cell RNA and protein measurements shown in Figure 7F**-7I** and **S7E**.

## 2 Supplementary Tables

Table S1. Oligonucleotides, probes and siRNA sequences

Table S2. GSEA Enrichment results: Proliferative vs differentiated HepaRG

Table S3. Empirically defined pre-tRNAs

Table S4. Normalized count tables

Table S5. ***R*** analysis of precursor-mature pairs

Table S6. GSEA Enrichment results: hiHep MYC-OE vs GFP-OE

## 3 Supplementary Note S1. Mathematical modeling and simulation of precursor-to-mature dynamics

### 3.1 Mathematical model

tRNA accumulation can be represented using a standard two-compartment kinetic model (**Figure 3A**) describing the dynamics of precursor and mature RNAs:

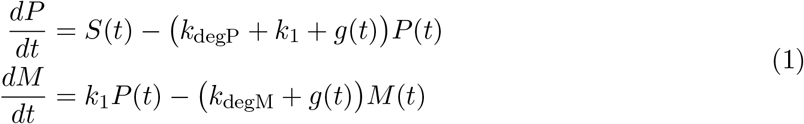

where *S*(*t*) denotes the transcription rate, *k*_1_ the processing rate, *k*_degP_ and *k*_degM_ the decay rates of precursor and mature tRNAs, respectively, and *g*(*t*) the dilution rate due to growth. At the onset of proliferation, *g*(*t*) transitions from zero to a positive value, reflecting re-entry into the cell cycle. In the simulations described below, this transition was approximated as an instantaneous switch at *t* = 0 to the experimentally measured proliferative growth rate *g* = 0.0184 h*^−^*^1^.

The precursor-to-mature ratio *R* = *P/M* is particularly suitable for RNA-seq data because it is independent of multiplicative scaling and robust to relative measurements.

Dividing the equation for mature tRNA dynamics (Equation (1)) by *M* (*t*) yields

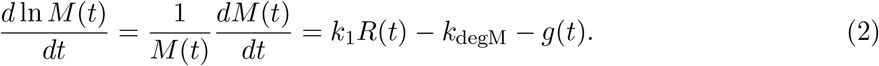

This relationship holds both at and away from steady state. Transcription and precursor degradation do not enter directly, as their effects on mature tRNA production are mediated through precursor abundance, captured by R(t). Thus, for a given rate of mature tRNA accumulation and unchanged mature tRNA decay and growth-associated dilution, a reduction in R implies an increase in the effective precursor-to-mature conversion rate constant.

Under constant growth and constant kinetic parameters (assuming *g*(*t*) *≈ g* after proliferation onset), the system reaches a steady state characterized by

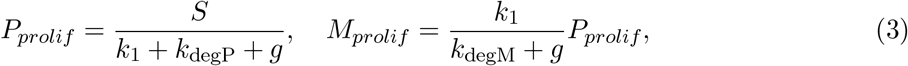

which implies a steady-state ratio

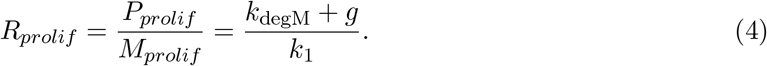

Importantly, the steady-state ratio is independent of the transcription rate *S* and depends solely on the processing and decay rates and the growth rate. Since *g >* 0 upon proliferation, the new steady-state satisfies

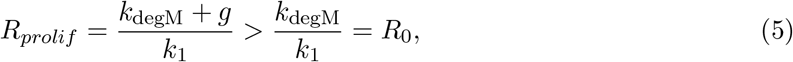

indicating that, if the system were allowed to reach its new steady-state, the precursor-to-mature ratio would eventually exceed its initial value (this prediction relies on post-transcriptional parameters remaining constant; deviations from this assumption are explored in simulations described below, see Section 3.2). This defines the null hypothesis: under transcription-only regulation, *R*(*t*) is expected to approach a value above *R*_0_*_h_* at long times.

### **3.2** Simulation-Based Exploration of Precursor-to-Mature Dynamics

The previous section established that, in absence of additional post-transcriptional events, the precursor-to-mature ratio *R* is higher for genes that have reached steady-state in the proliferative state compared to their steady-state levels in non-dividing, differentiated cells. Deviations from this expectation, i.e. *R_prolif_ < R*_0_*_h_*, would indicate that post-transcriptional parameters have changed, specifically precursor processing and/or mature tRNA decay relative to growth. However, in our experimental setting, because mature tRNAs can be highly stable, with half-lives exceeding tens of hours, the system may not reach the new steady-state within the first 48 h after the onset of re-programming. Transient deviations from *R_prolif_* are therefore expected even under transcription-only regulation, and *R*(*t*) may temporarily remain below *R*_0_*_h_* depending on precursor dynamics, initial conditions, and the timescale separation between rapid precursor turnover and slower mature tRNA dynamics. Even though slow adaptation to the new steady-state is unlikely to generate coherent decreases in ratios across the majority of genes, the central question becomes whether transient dynamics, under realistic transcriptional adaptation and growth, can lead to a systematic decrease of *R*_48_*_h_* below *R*_0_*_h_*. Alternatively, such behavior may require additional post-transcriptional regulation.

#### 3.2.1 Simulation Design and Parameterization

Because transcription rates were not directly observed and temporal sampling was sparse, we first assessed whether transcriptional regulation alone can account for observed *R*(*t*) ratios using numerical simulations of the classical two-compartment model (Equation (1)). The system of differential equations was solved numerically using odeint under a variety of transcriptional programs *S*(*t*) designed to capture biologically plausible adaptation kinetics and explore the feasible space of *R* dynamics. These simulations allowed us to characterize the expected behavior of *R*(*t*) at intermediate times, such as 48 h, when the system may not yet have reached the new steady-state *R_prolif_*.

Beyond purely transcription-driven dynamics, we also explored scenarios in which the processing rate *k*_1_(*t*) or mature tRNA decay rate *k*_degM_(*t*) changes over time. By comparing the resulting *R*(*t*) trajectories under different combinations of transcriptional and post-transcriptional regulation, we can delineate which aspects of the observed data can be explained by transcription alone and which require additional regulatory mechanisms.

##### Parameterization and Steady-State Initialization

Simulations were performed using a fixed set of kinetic parameters chosen to approximate experimentally observed regimes rather than to infer underlying rates. Parameter values were selected to reproduce the separation of timescales between precursor and mature tRNAs and the precursor-to-mature ratios observed in our data. Given the lack of direct measurements of processing and decay kinetics for individual tRNAs, these values define a plausible dynamic regime but are not uniquely constrained.

Specifically:

- The experimentally measured proliferative growth rate was fixed at *g* = 0.0184 h*^−^*^1^ (estimated in **Figure 1A**), corresponding to a doubling time DT = ln(2)*/g ≈* 37.8 h.
- The basal transcription rate was set to *S*_0_ = 100 (arbitrary units), which serves as a reference level for relative fold-changes in *S*(*t*).
- The processing rate from precursor to mature RNA was fixed at *k*_1_ = 4 h*^−^*^1^, chosen to reflect a relatively rapid processing time relative to mature RNA turnover.
- Precursor decay was modeled with a short half-life of approximately 20 min, corresponding to *k*_degP_ *≈* 2.08 h*^−^*^1^ in the absence of processing. Combined with the processing rate *k*_1_ = 4 h*^−^*^1^, this yields a shorter effective precursor lifetime similar to measured half-lives in this study (*∼*7 min).
- Two representative regimes of mature RNA stability were considered:

– a short-lived mature species with half-life DT*/*8 *≈* 4.7 h, and
– a long-lived species with half-life 2 *×* DT *≈* 75.5 h.

For each combination of parameters, the system was initialized in steady state, corresponding to the non-proliferative reference condition (*t* = 0):

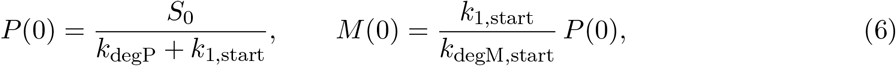

where *k*_1,start_ and *k*_degM,start_ denote the initial values of the time-dependent kinetic parameters for the corresponding regime. This ensures that at *t* = 0 the kinetic model matches the pre-stimulus steady state assumed in the analysis of real data.

##### Transcriptional Perturbation Families

To model transcriptional regulation following induction of proliferation, we defined a family of smooth, continuous transcriptional programs *S*(*t*). These programs were parameterized by biologically interpretable features:

- **Peak magnitude relative to** *S*_0_: from strong downregulation to strong upregulation (1*/*8*×* to 8 *× S*_0_).
- **Time to peak:** the time at which *S*(*t*) reaches its maximum or minimum relative to *S*_0_, sampled at 3, 6, 12, 24, or 48 h.
- **Plateau duration at peak:** how long transcription remains at the peak before relaxing, set to 0, 6 h, or *∞*.
- **Decay timescale:** exponential relaxation from the plateau toward the final transcription level, parameterized as either 1.5 *× t*_peak_ or 3 *× t*_peak_.
- **Final transcription level relative to** *S*_0_: the terminal level approached after the decay phase, set to 1*/*2*×*, 1*×*, or 2 *× S*_0_.

For each transcriptional program, *S*(*t*) was defined in three phases: (i) a normalized sigmoidal transition (rescaled to exactly match boundary conditions) from *S*_0_ to *S*_peak_ over the interval 0 *≤ t < t*_peak_, (ii) a plateau phase with *S*(*t*) = *S*_peak_ for *t*_peak_ *≤ t < t*_peak_ + *t*_plateau_, and (iii) an exponential relaxation toward *S*_end_ for *t ≥ t*_peak_ + *t*_plateau_ with time constant *t*_decay_. Upward and downward modulations were implemented using the same normalized sigmoidal logic, with *S*_peak_ *> S*_0_ or *S*_peak_ *< S*_0_, respectively, to ensure continuity and matching boundary conditions. For the analysis shown in **Figure 3I**, the timing of transcriptional relaxation was defined as *t*_peak_ + *t*_plateau_ + *t*_decay_, corresponding to one exponential relaxation time constant after the end of the plateau. At this time, transcription has completed approximately 63% of the transition from its peak toward its final level (*S*_end_). Programs with an indefinite plateau were assigned an infinite relaxation time.

##### Combinatorial grid and number of traces

For each of the six non-constant peak-magnitude conditions (1*/*8, 1*/*4, 1*/*2, 2, 4, and 8 *× S*_0_), transcriptional programs were generated by combining 5 values of *t*_peak_, 3 plateau durations, 2 decay timescales, and 3 final transcription levels, yielding 90 parameter combinations per condition and 540 in total. Some parameter combinations generated identical transcriptional trajectories within the 48 h simulation window (e.g., when the plateau extended beyond this period); these combinations were retained in the analysis. The unperturbed condition (*S*_peak_ = *S*_0_) was represented by three identical constant-transcription controls corresponding to the three nominal final-level settings. Thus, the grid comprised 543 parameter combinations per mature RNA stability regime, yielding 1,086 simulated trajectories across the two mature RNA half-life conditions.

**Table 1.** Parameters used to define transcriptional perturbation programs.

| **Parameter** | **Levels** | **Values** |
|---|---|---|
| Peak magnitude | 7 | 1/8, 1/4, 1/2, 1, 2, 4, 8 × *S*<sub>0</sub> |
| Time to peak | 5 | 3, 6, 12, 24, 48 h<sup>a</sup> |
| Plateau duration | 3 | 0, 6 h, ∞ <sup>a,b</sup> |
| Decay timescale | 2 | 1.5 × *t*<sub>peak</sub>, 3 × *t*<sub>peak</sub> <sup>a</sup> |
| Final transcription level | 3 | 1/2, 1, 2 × *S*<sub>0</sub> <sup>a</sup> |

<sup>a</sup> Applied only when peak magnitude ≠ 1 × *S*<sub>0</sub>. The three nominal final-level settings were retained for the unperturbed controls but had no effect on their transcription profiles.
<sup>b</sup> ∞ indicates that the plateau lasts until the end of the simulation.

#### 3.2.2 Simulation Execution and Coverage

##### Post-Transcriptional Regulation Regimes

In addition to transcriptional variation, we simulated a constant-kinetics reference regime together with a set of post-transcriptional kinetic perturbation regimes. These are defined by time-dependent changes in processing rate *k*_1_(*t*) and/or mature RNA decay rate *k*_degM_(*t*), modeled as exponential relaxations from an initial value toward a perturbed value with a fixed time constant *τ* :

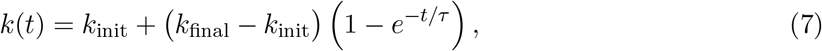

where *k*(*t*) denotes either *k*_1_(*t*) or *k*_degM_(*t*). The time constant was fixed to

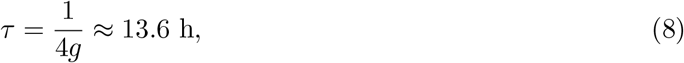

corresponding to approximately one third of the cell doubling time. This ensures that post-transcriptional modulation occurs on a timescale comparable to, but shorter than, the cell cycle. We specifically considered one constant-kinetics reference regime together with four post-transcriptional perturbation regimes summarized in Table 2.

**Table 2.** Post-transcriptional kinetic regimes simulated.

| Regime | Description |
|---|---|
| 1. Constant kinetics | *k*<sub>1</sub>(*t*) = *k*<sub>1</sub>, *k*<sub>degM</sub>(*t*) = *k*<sub>degM</sub> |
| 2. Decreased processing | *k*<sub>1</sub>(*t*) decreases to 0.25 × *k*<sub>1</sub>, *k*<sub>degM</sub>(*t*) constant |
| 3. Increased processing | *k*<sub>1</sub>(*t*) increases to 4 × *k*<sub>1</sub>, *k*<sub>degM</sub>(*t*) constant |
| 4. Increased stability | *k*<sub>degM</sub>(*t*) decreases to 0.25 × *k*<sub>degM</sub>, *k*<sub>1</sub>(*t*) constant |
| 5. Decreased stability | *k*<sub>degM</sub>(*t*) increases to 4 × *k*<sub>degM</sub>, *k*<sub>1</sub>(*t*) constant |

These regimes test whether post-transcriptional changes are required to reproduce the systematic decrease in *R*(*t*) observed experimentally.

##### Parameter values

The baseline values correspond to those used for simulations:

- *k*_1_ = 4 h*^−^*^1^, *k*_degP_ = ln(2)*/*0.33 h *≈* 2.1 h*^−^*^1^, and *k*_degM_ as described for short- and long-lived mature RNAs.
- Fold-changes applied to generate each regime: 1/4*×*, 1*×*, 4*×* relative to the baseline.
- Relaxation time constant: *τ* = 1*/*(4 *× g*).

##### Simulation coverage

The 543 transcriptional programs defined for each mature RNA stability regime were combined with all five kinetic regimes, yielding 1,086 trajectories per regime and exactly 5,430 simulated trajectories in total. This design ensures that each regime is systematically tested while keeping the computational load feasible.

#### 3.2.3 Numerical Integration and Time-Point Extraction

To facilitate comparison with the experimental sampling scheme, simulated trajectories were sub-sampled at *t* = 0, 12, 48 h. The precursor-to-mature ratio was also normalized relative to its steady-state value at *t* = 0:

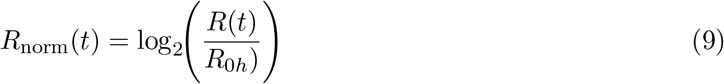

yielding quantities directly comparable to analogous metrics derived from the RNA-seq data.

##### Simulation Outputs and Usage

Simulation results were used to assess:

- whether *R*_48_*_h_ < R*_0_*_h_* can arise under transcriptional modulation only,
- how different combinations of transcriptional and post-transcriptional changes affect the temporal pattern of *R*(*t*),
- and how these patterns compare to the experimental observations.

This simulation ensemble forms the basis for Figure 3 and S3 in the main text.

#### 3.2.4 Implications for interpretation of *R* dynamics

Across the simulated parameter space, the classical two-compartment model with constant post-transcriptional parameters rarely reproduced decreases in *R*(*t*) at 48 h under transcription-only regulation, even when accounting for transient, non-steady-state dynamics. In contrast, such decreases arose more readily when precursor processing or mature RNA decay were allowed to vary over time.

These simulations therefore define the expected behavior of *R*(*t*) under the sampled transcription-only regimes and provide a reference framework for interpreting deviations observed experimentally. In particular, they support the inference that transcriptome-wide reductions in *R* are more consistent with coordinated post-transcriptional modulation than with transcriptional regulation alone.

